# Structural and biophysical correlation of anti-NANP antibodies with *in vivo* protection against *P. falciparum*

**DOI:** 10.1101/2020.07.18.210385

**Authors:** Tossapol Pholcharee, David Oyen, Yevel Flores-Garcia, Gonzalo Gonzalez-Paez, Zhen Han, Katherine L. Williams, Wayne Volkmuth, Daniel Emerling, Emily Locke, C. Richter King, Fidel Zavala, Ian A. Wilson

## Abstract

The most advanced *P. falciparum* circumsporozoite protein (PfCSP)-based malaria vaccine, RTS,S/AS01 (RTS,S), confers partial protection but with antibody titers that wane relatively rapidly, highlighting the need to elicit more potent and durable antibody responses. Here, we elucidate crystal structures, binding affinities and kinetics, and *in vivo* protection of eight anti-NANP antibodies (Abs) derived from an RTS,S phase 2a trial and encoded by three different heavy-chain germline genes. The structures reinforce the importance of homotypic Fab-Fab interactions in protective Abs and the overwhelmingly dominant preference for a germline-encoded aromatic residue for recognition of the NANP motif. A number of biophysical properties were analyzed and antibody affinity correlated best with protection in an *in vivo* mouse model, with the more potent antibodies also recognizing epitopes with repeating secondary structural motifs of type I β- and Asn pseudo 3_10_ turns. Such insights can be incorporated into design of more effective immunogens as well as antibodies for passive immunization.

## Introduction

Malaria is caused by unicellular eukaryotic *Plasmodium* parasites, and *P. falciparum* is responsible for most malaria morbidity and mortality. Despite significant progress over the past 20 years, resistance of mosquito vectors to pyrethroid (1) and the emergence of multidrug-resistant parasite strains (2) emphasize the need for new tools, including vaccines, to combat the disease. The most advanced malaria vaccine candidate to date is RTS,S/AS01 vaccine, which has completed phase 3 clinical trials in young African children and is currently undergoing a large scale pilot introduction in Malawi, Ghana, and Kenya to inform on a policy decision for broader use (3). The vaccine is based on PfCSP, which densely covers the surface of sporozoites and plays a critical role in the *P. falciparum* life cycle from the development of sporozoites in the mosquito midgut to liver-stage development in humans (4-7). The N-terminal domain of CSP includes a heparan sulfate binding site for hepatocyte adhesion (4), followed by the immunodominant central repeat region (8), and the C-terminal α-thrombospondin repeat (αTSR) domain that contains multiple T-cell epitopes (9). The repeat region in *P. falciparum* is composed of 1 NPDP, 3-5 NVDP, and 35-41 NANP repeats (10-13). In contrast, RTS,S contains only 19 NANP repeats and the αTSR domain, linked to the hepatitis B surface antigen protein (HBsAg), and was expressed recombinantly with soluble HBsAg to form a virus-like particle that is administered with the AS01 adjuvant (14). RTS,S displayed ∼40 % efficacy against clinical malaria disease over 4 years of follow-up in phase 3 clinical testing; vaccine efficacy is highest in the period immediately following immunization and declines coincident with decay of induced Ab titers to CSP (15-18). A similar vaccine candidate, R21, composed only of the same HBsAg-CSP fusion (i.e. without extra HBsAg), and formulated with Matrix-M adjuvant, has recently entered phase 2 clinical testing and is showing comparable efficacy levels in early clinical studies (19). Another candidate is the attenuated, whole-sporozoite-based PfSPZ vaccine (20), delivered by direct venous inoculation. It primarily aims to induce cellular immunity, and is thus associated with lower anti-CSP antibody titers compared to R21 and RTS,S; however vaccine efficacy in endemic field studies has been modest (21). These clinical studies highlight the need to improve current vaccines to induce either more durable protection and/or higher potency antibody responses.

Recently, many anti-CSP antibodies have been characterized using structural and biophysical approaches and various functional assays, which have contributed to our growing insights into humoral immune responses against CSP. The junctional region, which corresponds to the amino-acid sequence of PfCSP between the N-term domain and the NANP repeats, contains NPDP and NVDP motifs and has been shown as a target for potent antibody responses (22, 23). However, recent evidence suggests that anti-junction mAbs can also cross-react with the NANP repeats, and their protective capacity can be correlated with their binding promiscuity to NANP (24, 25). Epitopes of anti-NANP antibodies typically contain 2-3 NPNA structural motifs, which can adopt local conformations of a type I β-turn and an Asn pseudo 3_10_ turn (22, 23, 26-30). Anti-NANP antibodies often utilize a germline-encoded Trp residue to interact with the Pro or Asn in the NPNA turns (29). Additionally, some of these Abs exhibit unusual homotypic inter-Fab contacts when they simultaneously recognize the adjacent repeating epitopes on the NANP region of CSP (26, 27). Despite these structural and biophysical advances, the implications for antibody and immune responses, especially which properties or structural features correlate with *in vivo* protection by anti-NANP antibodies, are still not resolved.

Here, we characterized eight monoclonal antibodies (mAb) derived from protected volunteers who participated in a phase 2a clinical trial of RTS,S/AS01 (31) with a delayed fractional dosing regimen, and compared them with three previously published mAbs derived from the same clinical trial (28, 29). Our data suggest a correlation between antibody affinity, driven by the off rate, and *in vivo* protection, which could serve as an important basis for subsequent characterization and engineering of anti-NANP mAbs. Two antibodies exhibit homotypic Fab-Fab interactions, which increase avidity to the repeat peptides as found in previous studies, and may have implications for Ab responses against CSP as an unusual type of antigen (26, 27). Co-crystal structures also reveal conserved and convergent use of aromatic residues for interaction with the NANP repeat region. Furthermore, we observed that binding to an extended conformation of the NANP repeats, which lacks any secondary structural motifs, may not be optimal for stable antibody interaction and could contribute to low affinity to CSP and, subsequently, poor protection. On the other hand, potent and high affinity antibodies all recognize epitopes containing a type I β-turn and/or Asn pseudo 3_10_ turn, which are the two structural motifs observed as a repeating unit on a soluble recombinant shorter version of CSP (rsCSP), which can adopt an unusual, long-range, extended spiral conformation (27). Altogether, these comprehensive characterizations of anti-NANP antibodies enhance our understanding of human humoral immune responses against CSP and provide a strong foundation for the design of next-generation malaria vaccines.

## Results

### Many anti-NANP antibodies examined in this study show potent functional activity *in vivo*

The antibodies in this study were derived from protected volunteers in a phase 2a trial of RTS,S/AS01 seven days after the third fractional dose, as previously reported (27-29). Antibodies were selected from among expanded sequence families with a focus on prevalent *IGHV* families. Eight monoclonal antibodies (mAbs) were investigated that were encoded by 3 different immunoglobulin heavy variable (*IGHV*) genes and compared to previous mAbs 311, 317, and 397 from this set that include two other *IGHV* genes (27-29). Experiments to evaluate *in vivo* protection were conducted for two panels of antibodies using two mouse models that assess parasite liver burden load and blood-stage parasitemia (32, 33). The first panel contained mAbs derived from the *IGHV3-33* gene (mAb239, 337, 356, 364, and 395), and the second consisted of mAbs encoded by *IGHV3-49* (mAbs 224, 399), *IGHV1-2* (mAb366), *IGHV3-15* (mAb397), and *IGHV3-30* (mAb317) germline genes, with mAb311 (from *IGHV3-33*) as a control across both experiments. To assess reduction of parasite liver burden load, mice (*N = 5*) received 100 µg mAb by intravenous injection (IV) and, after 16 hrs, were challenged with chimeric sporozoites (*P. berghei* sporozoites expressing full-length *P. falciparum* CSP) (32) (Fig. 1A). All antibodies significantly reduce parasite burden (except for mAb395 (*p < 0*.*05)*; *p < 0*.*01* for other mAbs, Mann-Whitney U test) with most mAbs inhibiting parasite development by at least 93% (Fig. 1A). However, in the first panel, mAb395 performs significantly worse and mAb337 shows less *in vivo* protection compared to the others (*p < 0*.*01*, Mann-Whitney U test) (Table S1). In the second panel, mAbs 366 and 397 also display weaker protection, which is significantly less than mAbs 224 and 399 (*p < 0*.*05*, Mann-Whitney U test) (Table S1).

**Fig. 1.**
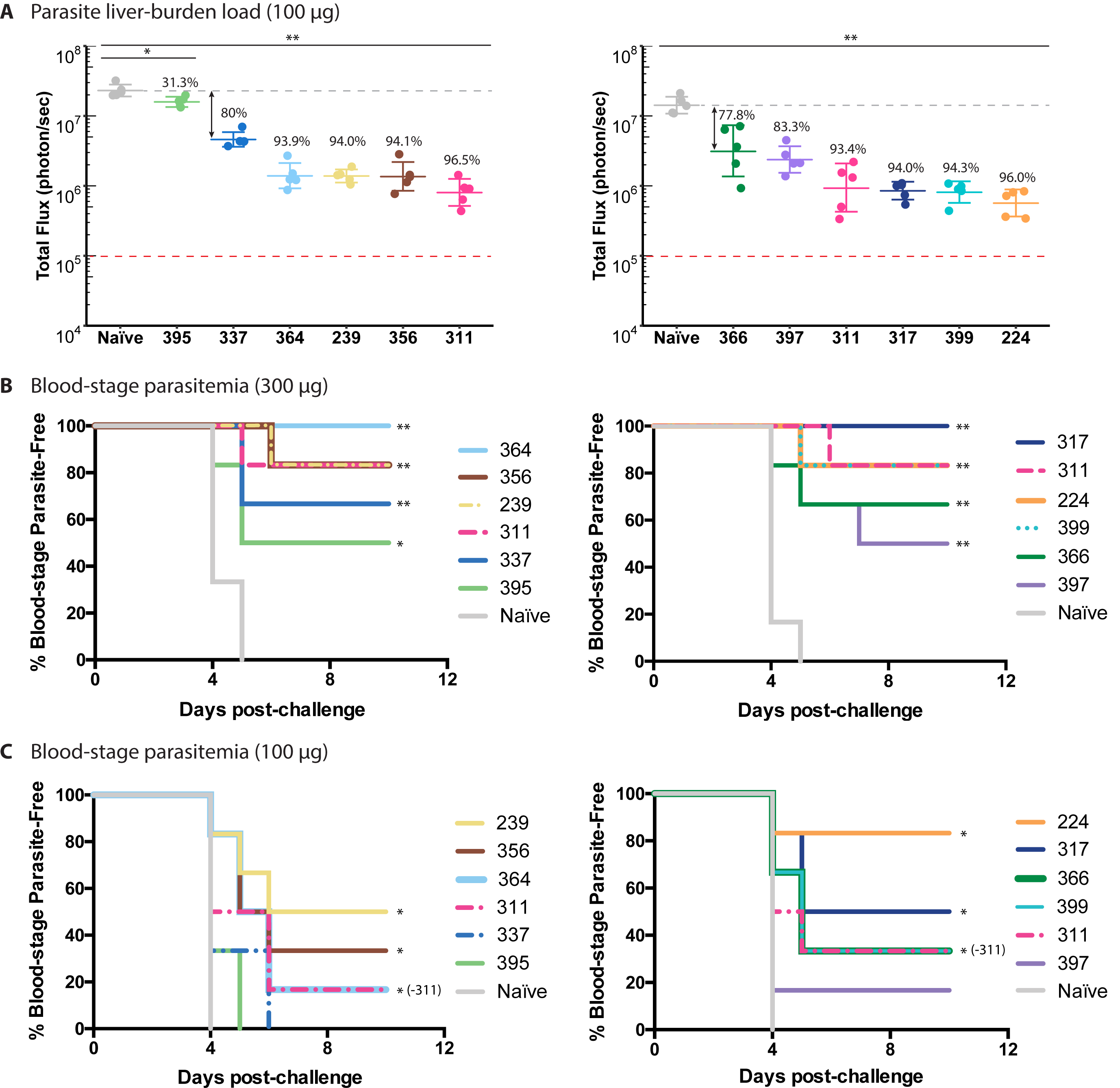
Assessment of antibody *in vivo* protection. **A)** Parasite liver burden load was measured by bioluminescence of *P. berghei* sporozoites expressing luciferase-conjugated *Pf*CSP after passive transfer of 100 µg of antibody in C57Bl/6 mice (*N = 5*). The numbers indicate the percent inhibition of the mean parasite burden relative to that of naïve control mice (i.e. % inhibition). A Mann-Whitney U test was used; \**p < 0*.*05*, and *\*\*p < 0*.*01*. Red dashed lines indicate the baseline signal from naïve non-infected mice treated with D-luciferin as established previously (32). Parasite-free mice after passive immunization with **B)** 100 µg or **C)** 300 µg of the indicated antibodies before challenge with bites of infected mosquitoes. A log-rank test was used; \**p < 0*.*05* (excluding mAb311 in **C**), and *\*\*p < 0*.*01* (*N = 6* per group).

To further validate the liver burden results, mAbs (100 µg or 300 µg) were passively transferred by IV to mice (*N = 6*) 16 hrs before exposure to bites of infected mosquitoes (Fig. 1B, C). At 300 µg mAbs, all antibodies from both panels protect at least 50% of the mice from blood-stage infection compared to the naïve control group (*p < 0*.*05*, log-rank test). Nonetheless, all mAbs, except for those that confer sterile protection, have overlapping confidence intervals (not shown), indicating insufficient statistical power (*N = 6*) to distinguish between these mAbs (Fig. 1B). For 100 µg mAbs, only mAb239, 356 and 364 from the first panel, and mAb224, 399, 366, and 317 from the second panel exhibit protection that is significantly greater than the naïve control mice (*p < 0*.*05*, log-rank test) (Fig. 1C).

### Antibody affinity substantially increases with homotypic Fab-Fab interactions

Antibody binding affinities were measured using isothermal titration calorimetry (ITC) against both short and long NANP repeat peptides to capture potential increases in apparent affinity due to avidity effects through homotypic Fab-Fab interactions, as observed previously for certain Fabs (26, 27). Despite conventional use of the term “NANP” repeats, our previous studies have consistently shown that anti-NANP antibody epitopes are typically composed of two to three “NPNA” structural motifs (27-29). Since structural characterization of mAb311 suggests that Fabs derived from the *IGHV3-33* gene have a minimal epitope of two NPNA repeats (27, 28), binding of mAbs 311, 239, 337, 356, 364, and 395 was measured against NPNA_2_, NPNA_4_, and NPNA_6_ peptides, whereas Fabs derived from other germline genes were tested with NPNA_3_ or NPNA_4_ as short peptides, and NPNA_6_ and NPNA_8_ as long peptides. Binding affinities for mAbs 397 and 317 were taken from our previous data (28, 29).

While some Fabs have strong binding (nM range) against short NPNA peptides, most Fabs tested here start from low affinity (µM range) that is dramatically increased against longer NPNA peptides, as indicated by the fold-change of the dissociation constants (K_d_) (Table 1, Figs. S1 and S2, and Table S2). For example, mAb356 recognizes NPNA_2_ with a µM K_d_ but binds NPNA_6_ in the nM range, resulting in an ∼300-fold K_d_ change (Table 1). Overall, most mAbs bind strongly to the longer NPNA peptides, except for mAb395 and 366, which remain at µM affinity (Table 1). Multiple copies of Fab311, with an 8-fold mean Kd change between NPNA_2_ and NPNA_6_, simultaneously recognize rsCSP (NPDP/NVDP/NANP repeat ratio of 1/3/19 instead of 1/4/38 for the *P. falciparum* 3D7 strain) and exhibit homotypic inter-Fab contacts (27). Therefore, mAbs 239, 337, 356, and 399 with higher fold-changes should also exhibit Fab-Fab interactions on binding multiple epitopes on the CSP NANP repeat region (Table 1).

**Table 1.**
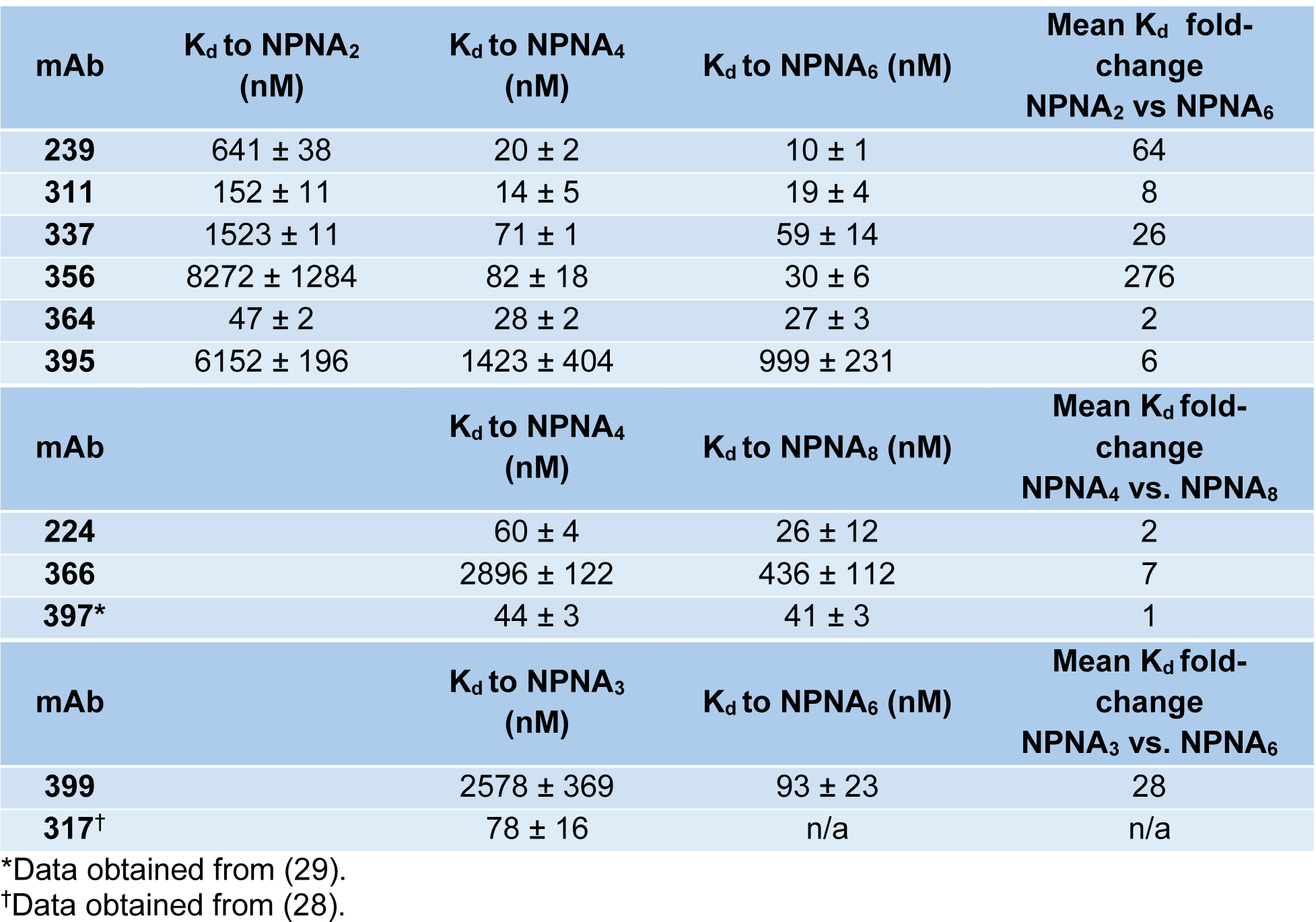
Dissociation constants of antibody Fabs and fold-changes in affinity obtained from ITC.

| mAb | K<sub>d</sub> to NPNA<sub>2</sub><br>(nM) | K<sub>d</sub> to NPNA<sub>4</sub><br>(nM) | K<sub>d</sub> to NPNA<sub>6</sub> (nM) | Mean K<sub>d</sub> fold-change<br>NPNA<sub>2</sub> vs NPNA<sub>6</sub> |
| --- | --- | --- | --- | --- |
| 239 | 641 ± 38 | 20 ± 2 | 10 ± 1 | 64 |
| 311 | 152 ± 11 | 14 ± 5 | 19 ± 4 | 8 |
| 337 | 1523 ± 11 | 71 ± 1 | 59 ± 14 | 26 |
| 356 | 8272 ± 1284 | 82 ± 18 | 30 ± 6 | 276 |
| 364 | 47 ± 2 | 28 ± 2 | 27 ± 3 | 2 |
| 395 | 6152 ± 196 | 1423 ± 404 | 999 ± 231 | 6 |
| mAb |  | K<sub>d</sub> to NPNA<sub>4</sub><br>(nM) | K<sub>d</sub> to NPNA<sub>8</sub> (nM) | Mean K<sub>d</sub> fold-change<br>NPNA<sub>4</sub> vs. NPNA<sub>8</sub> |
| 224 |  | 60 ± 4 | 26 ± 12 | 2 |
| 366 |  | 2896 ± 122 | 436 ± 112 | 7 |
| 397* |  | 44 ± 3 | 41 ± 3 | 1 |
| mAb |  | K<sub>d</sub> to NPNA<sub>3</sub><br>(nM) | K<sub>d</sub> to NPNA<sub>6</sub> (nM) | Mean K<sub>d</sub> fold-change<br>NPNA<sub>3</sub> vs. NPNA<sub>6</sub> |
| 399 |  | 2578 ± 369 | 93 ± 23 | 28 |
| 317<sup>†</sup> |  | 78 ± 16 | n/a | n/a |

\*Data obtained from (29).
<sup>†</sup>Data obtained from (28).

The presence of homotypic Fab-Fab interactions in mAbs 239 and 399 was validated in the crystal structures of Fab239-NPNA_4_ and Fab399-NPNA_6_ complexes (Fig. 2, Table S3). Two copies of Fab239 form side-to-side inter-Fab contacts, which are mediated mostly by the heavy chain (Fig. 2A-C). These Fab-Fab interactions are asymmetric where complementarity-determining region (CDR) H1 and H3 in one Fab interact mainly with CDR H2 and L3 in the adjacent Fab (Fig. 2C, G), and which is strikingly similar to what is observed in the cryo-EM structure of Fab311 with rsCSP (27). The interaction of ^H^Asp^31^ from CDR H1 with ^H^Glu^64^ from CDR H2 is also conserved in the Fab-Fab interfaces of Fabs 239 and 311 (Fig. 2C). Therefore, these similarities suggest that Fab239 may also be capable of forming a long-range spiral structure, in which multiple copies of the Fab simultaneously bind to rsCSP (27). Fab399 exhibits a different type of Fab-Fab interaction with a head-to-head configuration, resembling that of Fab1210 (26) (Fig. 2D, E), but the inter-Fab contacts here are perfectly symmetric, more extensive, and mediated almost entirely by the heavy chain. ^H^Asp^31^ in CDR H1 now forms a different network of hydrogen bonds and a salt bridge with ^H^Tyr^32^ and ^H^Arg^94^ in the adjacent Fab (Fig. 2F, G). Unlike Fabs 239 or 311, Fab399 is not likely to form a similar spiral conformation with rsCSP (27), because such a structure would not accommodate the symmetric, head-to-head, inter-Fab interactions. Despite the different CDR contributions in the homotypic interactions of Fabs 239 and 399, both antibodies similarly use CDR H1, H2, H3 and L3 for interaction with the NANP peptide (Fig. S3).

**Fig 2.**
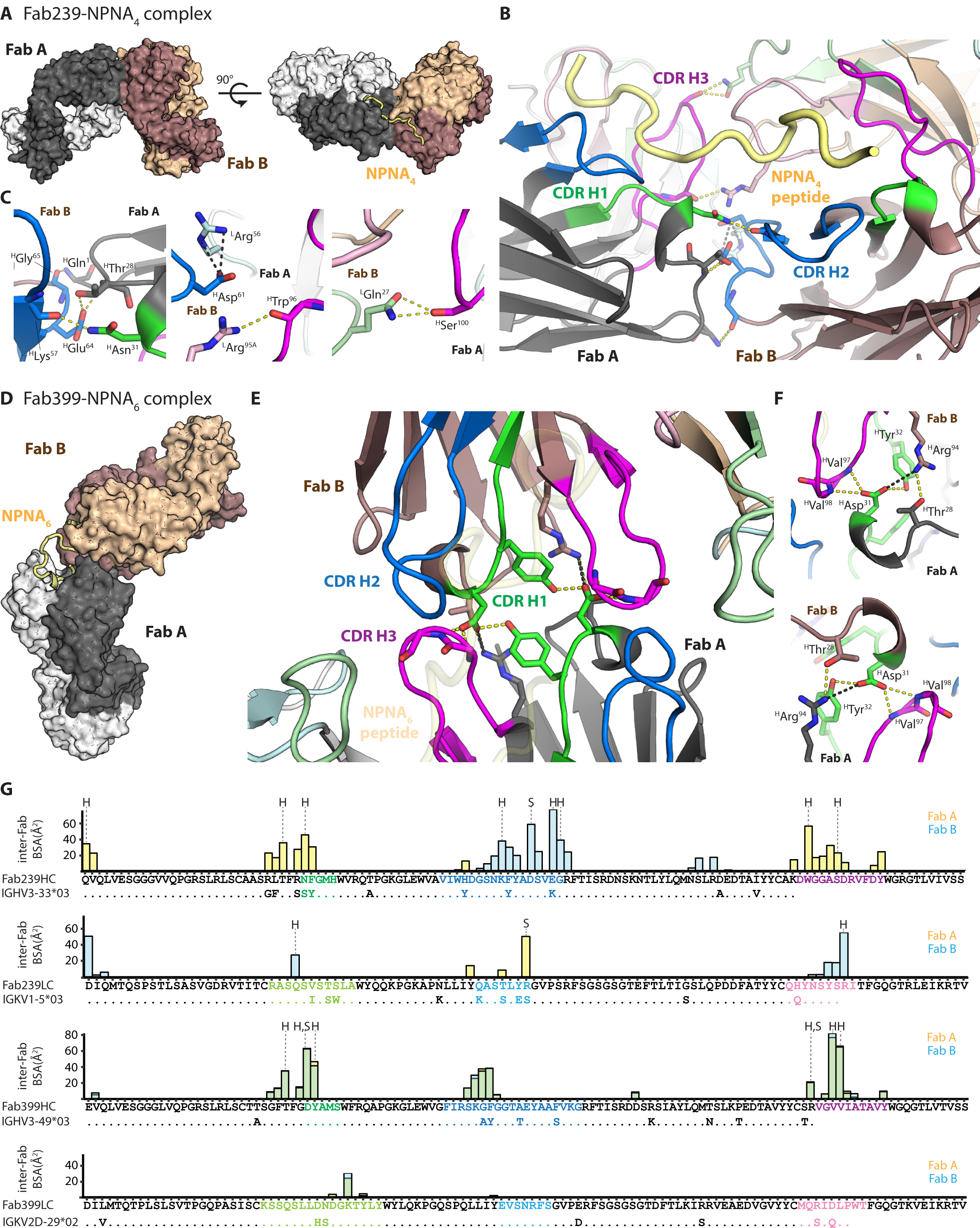
Homotypic Fab-Fab interactions in Fab239- and Fab399-peptide complexes. **A)** Two copies of Fab239 (Fab A, black: heavy chain, white: light chain. Fab B, dark brown: heavy chain, tan: light chain) in the crystal structure are shown as surfaces with the NPNA_4_ peptide represented as a yellow tube. **B)** and **C)** Interactions between two Fabs (A and B) that simultaneously recognize an NPNA_4_ peptide. Hydrogen bonds (yellow dashes) and salt bridges (black dashes) are highlighted. The Fabs are shown as cartoon representations with the side chains of interacting residues represented as sticks. CDRs as defined by Kabat are colored green, blue, magenta, light green, light cyan, and pink for CDR H1, H2, H3, L1, L2, and L3, respectively. **D)** Surface representation of homotypic, head-to-head interactions of two copies of Fab399 in the crystal structure with NPNA_6_ peptide (yellow tube). **E)** and **F)** Contacts between two Fabs that simultaneously recognize the NPNA_6_ peptide (transparent yellow tube). All coloring schemes and representations are as for the Fab239-NPNA_4_ complex. **G)** Individual residue contributions to the BSA of inter-Fab interactions are shown in a bar plot for the heavy and light chains of Fab239 and Fab399. The yellow and blue bars indicate the BSA on Fab A and Fab B (defined as in previous panels), respectively, while the green bars show the overlap of both. The CDRs are colored as in previous panels. Additionally, the alignment between the Fab heavy/light chain sequences and germline *IGHV* and *IGKV* gene sequences are shown to display somatically mutated residues. “H” and “S” mark residues that are engaged in hydrogen bonds and salt bridges, respectively.

### Antibody *in vivo* protection correlates with Fab binding affinity

To approximate the binding affinity and kinetics of these Fabs to full-length CSP, bio-layer interferometry (BLI) was performed using rsCSP (Table 2, Fig. S4). The overall K_d_ values with BLI are similar to those obtained from ITC, with mAbs 395 and 366 displaying the lowest affinity in the µM range (Tables 1 and 2). Most mAbs with high affinity exhibit potent functional activity in the mouse model of parasite liver burden load, except for mAbs 337 and 397, which perform worse than expected. The low affinity mAbs 366 and 395 perform less well, as anticipated (Table 2). Even although only 11 antibodies are examined in this study, the percent inhibition of parasite load, normalized across the two antibody panels, exhibits a correlation with Fab affinity to long NPNA peptides by ITC and to rsCSP by BLI with R^2^ of 0.8801 and 0.9257, respectively (Table 2 and Fig. 3). For cross-validation of the linear regression, bootstrapping was performed to generate 1,000 models, which yielded an average R^2^ (R^2^-boot) of 0.6777 and 0.7763 for affinity against both long peptide and rsCSP vs. normalized percent inhibition, respectively, which are lower than the R^2^ of the original models, but still suggest a linear correlation (Fig. 3). All antibodies have equivalent association rate constants (k_on_) in the order of 104 M-1s-^1^, which is not correlated with *in vivo* protection from the liver burden assay (Table 2 and Fig. 3). On the other hand, the most potent blocking antibodies display comparable dissociation rate constants (k_off_) in the range of 10^−^ _3_ s^-1^. The less potent mAbs 366 and 395 have faster k_off_ of ∼10^−2^ and ∼10^−1^ s^-1^, respectively (Table 2). Thus, the k_off_ component of the K_d_ correlates well with the normalized percent inhibition from the liver burden assay (R^2^ = 0.9230, R^2^-boot = 0.7633) (Table 2 and Fig. 3). Antibody thermal stability (T_m_) was also examined but showed no correlation with antibody activity as measured by liver burden load (Table 2). Thus, the outlier mAbs 337 and 397 with high affinity but poor protection could result from poor pharmacokinetics, e.g. durability and clearance in mice, aggregation in mouse serum, or cross-reactivity with mouse antigens etc.

**Table 2.** Dissociation and rate constants of antibody Fabs to rsCSP obtained from bio-layer interferometry (BLI) displayed with mean dissociation constants measured to the longest peptide tested by isothermal titration calorimetry (ITC), mean % inhibition of parasite burden studies, % blood-stage parasite free mice from parasitemia studies, total paratope buried surface area (BSA), number of hydrogen bonds between paratope and epitope, and melting temperature (T_m_) from differential titration calorimetry.

| mAb | $K_d$ (nM)<br>(ITC,<br>NPNA<sub>6/8</sub>) | $K_d$ (nM)<br>(BLI,<br>rsCSP) | $k_{on}$ (1/M·s) | $k_{off}$ (1/s) | LB<br>study 1 | LB<br>study 2 | Parasit.<br>study 1 | Parasit.<br>study 2 | BSA<br>(Å<sup>2</sup>) | #<br>H-bond | $T_m$<br>(°C) | Epitope<br>secondary<br>structure | |
| --- | --- | --- | --- | --- | --- | --- | --- | --- | --- | --- | --- | --- | --- |
|  |  |  |  |  |  |  |  |  |  |  |  | β-turn | 3<sub>10</sub><br>turn |
| 317 | 78 | 9.3 | $2.32 \times 10^4$ | $2.15 \times 10^{-4}$ | | 94.0 | 100.0% | | 519 | 7 | 73.3 | 3 | - |
| 224 | 26 | 16 | $5.69 \times 10^4$ | $8.97 \times 10^{-4}$ | | 96.0 | 83.3% | | 617 | 10 | 68.9 | 3 | - |
| 399 | 93 | 26 | $6.77 \times 10^4$ | $1.78 \times 10^{-3}$ | | 94.3 | 83.3% | | 492 | 7 | 82.0 | 2 | 1 |
| 311 | 19 | 31 | $3.83 \times 10^4$ | $1.19 \times 10^{-3}$ | 96.5 | 93.4 | 83.3% | 83.3% | 473 | 7 | 71.3 | 1 | 1 |
| 364 | 27 | 32 | $4.53 \times 10^4$ | $1.45 \times 10^{-3}$ | 93.9 | | | 100.0% | 443 | 6 | 72.1 | 1 | 1 |
| 356 | 30 | 42 | $3.89 \times 10^4$ | $1.62 \times 10^{-3}$ | 94.1 | | | 83.3% | 572 | 6 | 74.4 | 1 | 1 |
| 239 | 10 | 187 | $2.93 \times 10^4$ | $5.48 \times 10^{-3}$ | 94.0 | | | 83.3% | 550 | 8 | 71.8 | 1 | 1 |
| 397 | 41 | 63 | $4.92 \times 10^4$ | $3.10 \times 10^{-3}$ | | 83.3 | 50.0% | | 593 | 10 | 74.4 | 1 | 1 |
| 337 | 59 | 97 | $5.41 \times 10^4$ | $5.22 \times 10^{-3}$ | 80 | | | 66.6% | N/A | N/A | 85.2 | N/A | N/A |
| 366 | 436 | 1076 | $2.57 \times 10^4$ | $2.76 \times 10^{-2}$ | | 77.8 | 66.6% | | 576 | 13 | 81.3 | - | - |
| 395 | 999 | 4902 | $3.96 \times 10^4$ | $1.94 \times 10^{-1}$ | 31.3 | | | 50.0% | 420 | 7 | 80.3 | 1 | - |

**Fig 3.**
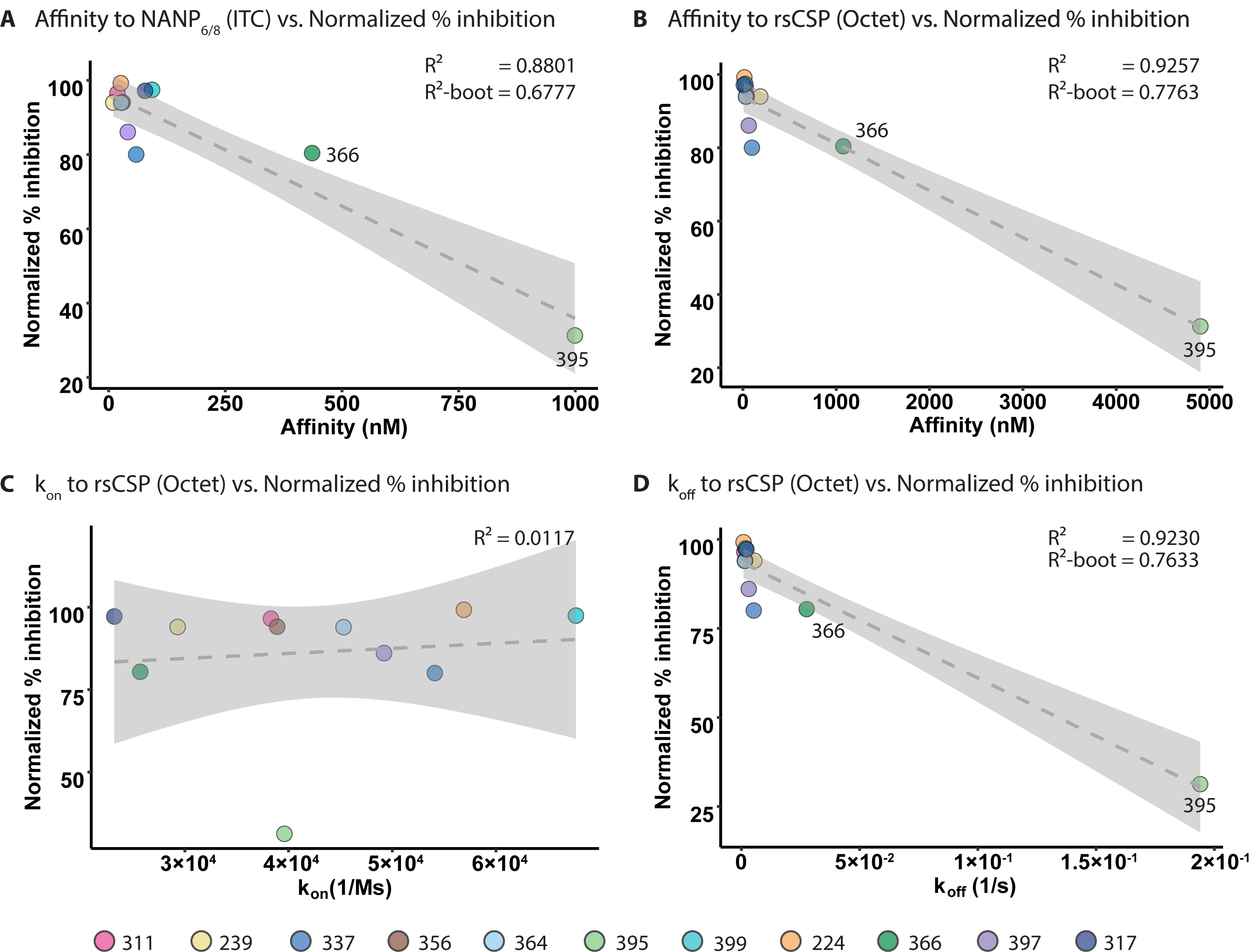
Correlation of affinity and dissociation constants with normalized parasite burden. The linear regression graph plots the % inhibition of the parasite liver burden load, normalized across the two mAb panels, against: **A)** Fab affinity as the dissociation constant (K_d_) measured to the NPNA_6_ or NPNA_8_ peptide using isothermal titration calorimetry; **B)** K_d_, **C)** association rate constant (k_on_), and **D)** dissociation rate constant (k_off_) measured against rsCSP using bio-layer interferometry. The dashed line indicates the fitted linear regression model with 95% confidence interval shaded in grey. The R^2^ value for each model is displayed, together with the average R^2^ from 1,000 models from bootstrapping (R^2^-boot). Data points for each mAb are colored as shown.

### Crystal structures of Fab-peptide complexes reveal conserved interaction motifs and structural basis of affinity and protection

To gain a better understanding of the molecular recognition of the antibodies to the NANP repeats and what structural features might relate to low binding affinity and poor protection, we co-crystalized Fabs 239, 356, 364, 395, 224, 399, and 366 with repeat peptides (NPNA_2,3,4, or 6_), and compared with our previous crystal structures of Fabs 311, 317, and 397 (28, 29).

Except for Fab395, the epitopes of the *IGHV3-33* Fabs share a similar conformation, which includes a type I β-turn and an Asn pseudo 3_10_ turn formed by the first and the second NPNA motifs, respectively. Although the Fab395 epitope is shorter (^1^NPNANP^6^), the first NPNA motif also adopts a type I β-turn (Fig. 4B, Table S4). The NANP peptides in the *IGHV3-33* Fabs interact mainly with the heavy chain as indicated by its dominant contribution to the buried surface area (BSA); the light-chain interactions are mostly mediated through CDR L3 (Fig. 4A, B, and Table S5). The *IGHV3-33* Fabs all have similar conformations of CDR H1 and H2, which contribute to conserved interactions with their epitopes, with Fab395 having a unique disulfide bond between ^H^Cys^29^ in the framework region and ^H^Cys^32^ in CDR H1 (Kabat nomenclature) (Fig. 4C, and Table S6). Notably, the main chain of residues in CDR H1 form a conserved hydrogen bonding network with the NANP peptide. Conservation of van der Waals interactions are mediated by aromatic residues H32/33 in CDR H1, and H52 and H58 in CDR H2 (Fig. 4C, and Table S6). While Tyr in H32/33 and H58 can evolve to other aromatic residues, ^H^Trp^52^ is strictly conserved in our antibody panel. Two of these residues, H52 and H58, interact specifically with Pro in the Asn pseudo 3_10_ turn and type I β-turn respectively (Fig. 4C, and Table S6). Although Fab395 displays a seemingly different epitope compared to other *IGHV3-33* Fabs in this panel, the type I β-turn in the Fab395 epitope is positioned in the pocket where the Asn pseudo 3_10_ turn is accommodated in other Fabs, such as Fab311 (Fig. 4B). The Pro in the type I β-turn now interacts with ^H^Trp^52^ and the following Pro contacts ^L^Tyr^98^. The presence of the bulky ^L^Tyr^94^ in CDR L3 of Fab395 may prevent binding of the repeat peptide to the usual pocket for the type I β-turn as in other *IGHV3-33* Fabs (Fig 4B). Consequently, the binding pocket of Fab395 is considerably smaller and may contribute to its low binding affinity and poor protection.

**Fig 4.**
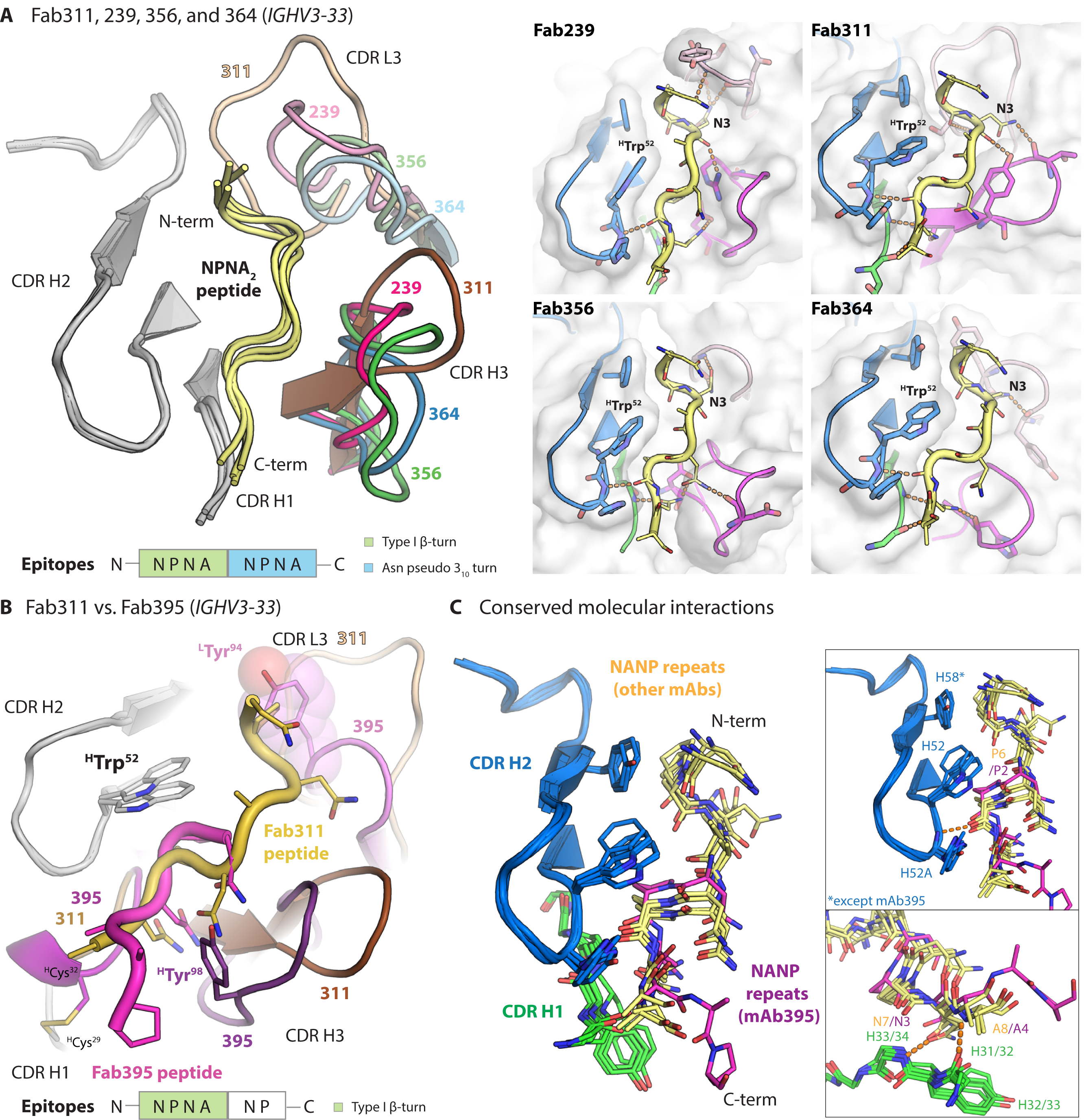
Crystal structure of *IGHV3-33* Fabs. **A)** Structures of Fab239, 311, 356, and 364 in complex with NPNA_2_ peptide (yellow) are shown in cartoon representation and aligned based on CDR H2. Only the CDRs involving in peptide binding are shown with CDR H1 and H2 colored in grey and CDR H3 and L3 colored for the different antibodies as indicated. A schematic of the epitope structural motifs is also indicated below. Close-up views of the paratopes are also displayed with the Fabs as cartoons embedded in their surface representation. CDR H1, H2, H3, and L3 are colored green, blue, magenta, and pink, respectively, and the peptides are shown as yellow tubes with side chains in stick representation. Antibody side chains engaging in hydrogen bonds (orange dashes) and key interacting aromatic residues are also shown as sticks. **B)** The paratopes of Fab395 aligned to that of Fab311 based on CDR H2 (grey) are displayed as cartoons with their CDRs colored as shown, with the schematic of the Fab395 epitope structural motif also indicated below. The side chains of ^H^Trp^52, H^Tyr^98^, and ^L^Tyr^94, H^Cys^29^, and ^H^Cys^32^ are highlighted as sticks (also with a surface representation for ^L^Tyr^94^). The peptides are shown as tubes with side chains as sticks and colored as indicated. **C)** Side chains of residues involving in conserved molecular interactions from CDR H1 (green) and H2 (blue) and the peptides are shown as sticks. The peptide bound to Fab395 is colored magenta, whereas others are in yellow. Hydrogen bonds are displayed as orange dashes.

The crystal structures of Fab224 and Fab399, derived from the same variable heavy chain *IGHV3-49* gene, also display similarities in their recognition of the NANP repeats (Fig. 5A, B, and Table S3 and S4). The epitopes are mainly composed of three NPNA motifs (Fab399 only has NPN in the last motif) with the first two NPNA motifs adopting a type I β-turn and the last motif exhibiting a type I β-turn in Fab224 and an Asn pseudo 3_10_ turn in Fab399 (Fig. 5A, B). The interactions of Fabs 224 and 399 with the NANP peptides are mediated mainly through the heavy chain (Fig. 5A, B and Table S5), which resembles those observed in the *IGHV3-33* antibodies (Fig. 4A, B). The binding pocket formed by the *IGHV3-49* heavy chains in these two Fabs both recognize the type I β-turn through nearly identical molecular interactions, including a CH-π interaction between the germline-encoded ^H^Phe^50^ and Pro in the NPNA repeat (Fig. 5A, B). Other shared interactions involve hydrogen bonds and van der Waals interactions mediated by ^H^Arg^52^ and ^H^Tyr^53^, or the somatically mutated ^H^Phe^53^ in Fab399, and hydrogen bonds from the CDR H3 backbone to the NANP peptide (Fig. 5A, B). Fabs 224 and 399 both have a Trp in CDR H3 (^L^Trp^96^) that hydrogen bonds with the backbone of the Ala preceding the conserved NPNA type I β-turn (Fig. 5A, B). However, their light chains are derived from different germline genes (Table S5). The Fab 399 epitope also shares striking similarities with that of a recently published, *IGHV3-49*-encoded Fab4493 in its crystal structure with the junctional peptide, GNPDPNANPN (24). The PDPNANPN core of the Fab4439 epitope displays a nearly identical conformation to the ANPNANPN residues in the Fab399 epitope, and both epitopes make similar interactions with ^H^Phe^50, H^Arg^52^, and ^L^Trp^96^ in the antibody paratopes (24).

**Fig 5.**
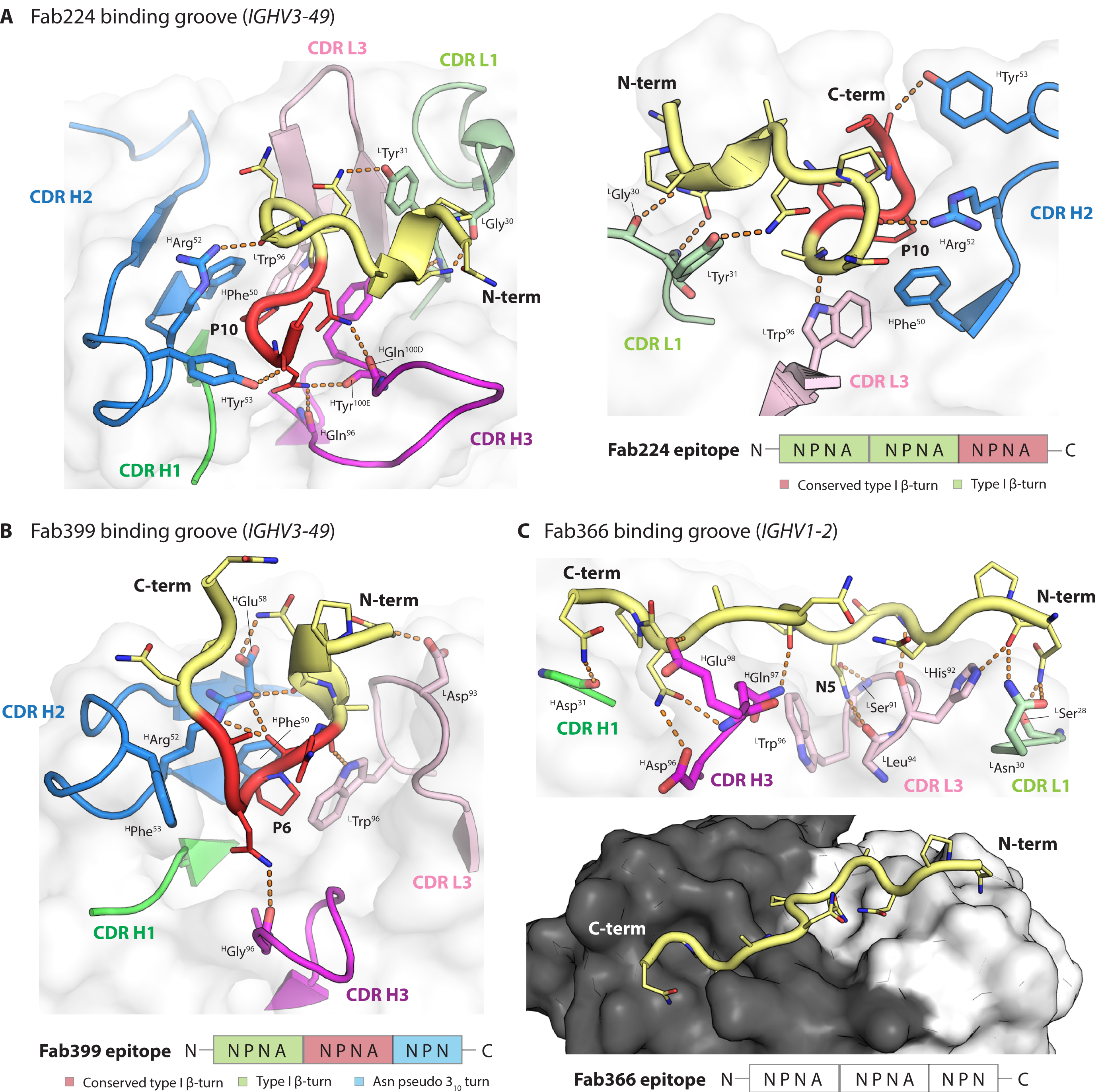
The paratopes of Fab224, 399, and 366. **A)** Crystal structure of Fab224 encoded by the *IGHV3-49* gene and its epitope are shown as cartoons with residues where the main chain and side chain engage in hydrogen bonds (orange dashes) shown as sticks. The Fab cartoon representation is embedded in a transparent white surface rendering. Only the CDRs involved in the binding groove are displayed and colored green, blue, magenta, light green, and pink for CDR H1, H2, H3, L1, and L3, respectively. A schematic of the epitope is also illustrated below. **B)** The binding groove of Fab399 encoded by the *IGHV3-49* gene. **C)** The paratope of Fab366 encoded by the *IGHV1-2* gene. All representations and coloring schemes for **B)** and **C)** are as in **A)**. In addition, a surface representation of Fab366 is shown, where the black and white surfaces represent the heavy and light chains, respectively.

We also determined the structure of Fab366 derived from the *IGHV1-2* gene, which has not been reported previously in anti-NANP antibodies. Its paratope is composed from CDR H1, H3, L1 and L3 without contribution from CDR H2, which is so dominant in Fabs from the *IGHV3-33* and *IGHV3-49* families (Fig. 5C, and Table S4). The epitope (from N-to C-terminus) spans a shallow binding groove from light to heavy chain and makes extensive hydrogen bonds with Fab residues. One notable interaction is between ^L^Trp^96^ and Asn5 in the peptide, which is similar to that observed in the Fab317- and Fab397-peptide complexes (28, 29) (Fig. 5C). Interestingly, the Fab366-bound peptide does not contain any secondary structural motifs (i.e. type I β-turn and Asn pseudo 3_10_ turn), but adopts a more extended conformation. In fact, the observation of a shallow Fab paratope recognizing extended NANP repeats resembles the epitope of Fab1450, which also has low affinity and poor *in vivo* protection (26) (Fig. 5C). Despite the extensive hydrogen-bonding network and BSA contribution within the range of the other Fabs analyzed here (Fig. 5C, and Table S5), Fab366 has a fast k_off,_ hence low affinity (Table 2), which seems attributable to a shallow binding groove, consistent with its similarity to Fab1450.

Another possible structural correlate of high affinity and better protection is the presence of local secondary conformations in the NANP epitopes, which are the type I β-turn and Asn pseudo 3_10_ turn with a hydrogen bond between the side chain of Asn (residue i) and backbone nitrogen of Asn (residue i+2) (Fig. 6). The peptides in potent and high affinity *IGHV3-33* antibodies (mAbs 239, 311, 356, and 364) contain both an NPNA type I β-turn and Asn pseudo 3_10_ turn (Table 2, and Fig. 6). Interestingly, the spiral rsCSP contains these two turns as a repeating unit, as observed in the cryo-EM structure with Fab311 (27). The epitopes of protective mAbs 317, 224, and 399 consist of up to 3 type I β-turns, but also display backbone-to-backbone H-bonds, resulting in a more compact conformation, compared to those with the *IGKV3-33* antibodies, and also higher affinity to rsCSP (Table 2, and Fig. 6). The mAb397 epitope exhibits both type I β- and Asn pseudo 3_10_ turns (Fig. 6), which correspond to the structural repeat in the long-range, curved conformation when multiple copies Fab397 bind to rsCSP (29). However, other factors such as antibody pharmacokinetics may account for the poor protection despite high binding affinity of mAb397. In contrast, the low affinity and weakly protective mAb395 has a shorter epitope that contains one type I β-turn, whereas the extended epitope of mAb366 is stabilized by only one H-bond between the side chain of Asn3 (i) and backbone oxygen of Asn5 (i+2), leading to a turn that is wider and more open than a type I β-turn (Fig. 6). Consequently, the extended epitopes of mAb366 when repeated in CSP may not be capable of adopting either the long-range curved or spiral conformations seen in mAbs 397 and 311, respectively.

**Fig 6.**
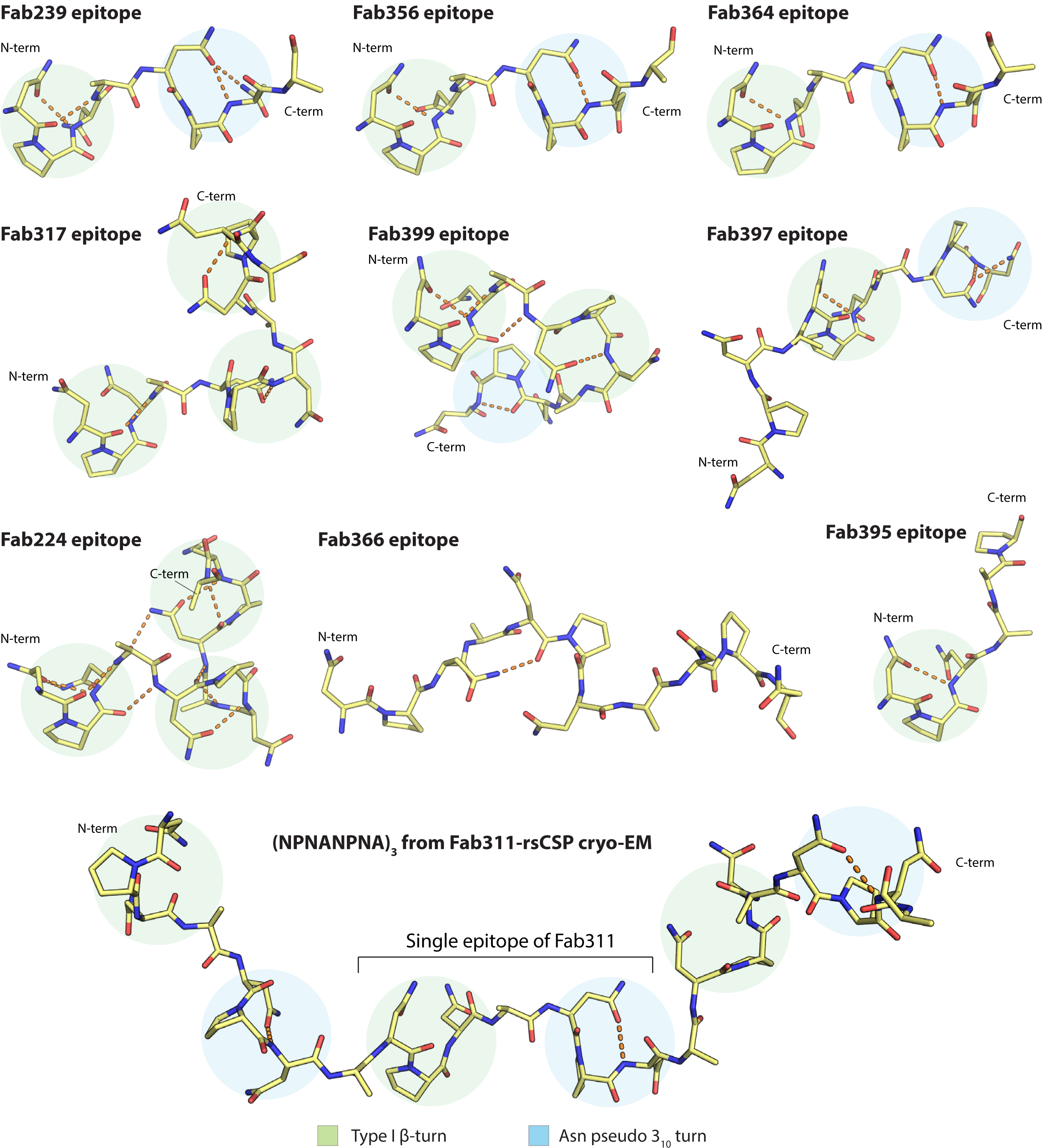
Summary of the epitope conformations for all antibodies analyzed in this study. The peptide epitopes are represented with yellow sticks, and the hydrogen bonds are shown in orange dashes. The type I β-turns and Asn pseudo 3_10_ turns are highlighted with green and blue circles. All peptides were aligned based on the first type I β-turn that is present, except for the epitope of Fab366 which was aligned to the overall epitope of Fab239. The epitope of Fab317 and Fab311 were obtained from the crystal and cryo-EM structures from previous studies (27, 28) (PDB ID: 6AXL and 6MB3, respectively).

## Discussion

This study strengthens and extends previous observations of recurring features of antibody recognition to the NANP repeats that pertain to their functional activity. Notably, we explored antibodies that were encoded by different sets of *IGHV* genes, the mechanisms by which different antibodies can achieve high Fab binding affinity, and the consequences for functional protection in liver burden and parasitemia models of infection. We also identified two additional antibodies, mAb239 (*IGHV3-33*) and mAb399 (*IGHV3-49*), that display homotypic inter-Fab contacts, while simultaneously recognizing NPNA repeat epitopes, as observed previously in mAbs 311 and 1210 (both from *IGHV3-33*) (26, 27). Fabs 239 and 311 exhibit an ‘asymmetric’, side-to-side Fab-Fab interaction with propensity to form a supramolecular, extended spiral structure with rsCSP, whereas Fab399 shows ‘symmetric’ inter-Fab contacts with a head-to-head configuration that may not be consistent with forming the same type of spiral architecture, and may be more similar to that observed in Fab1210 (26). We note that some of the key residues involved in homotypic Fab-Fab interactions are already present in the germline genes for both Fabs 239 and 399 (Fig. 2G). Indeed, most residues involved in inter-Fab contacts in Fab399 are germline-encoded, such as ^H^Thr^28, H^Asp^31, H^Tyr^32^, and ^H^Arg^94^ (Figs. 2F, G). Therefore, these two antibodies may exhibit a propensity for inter-Fab contacts, even prior to somatic hypermutation (SHM). A similar observation is found in rituximab (RTX) against CD20, a completely unrelated therapeutic antibody (34). Two copies of the Fab display homotypic interactions upon binding to their epitope on the CD20 dimer, resulting in a cross-linked circular supra-assembly of three RTX IgGs and three CD20 dimers (34). Intriguingly, all residues involved in the RTX Fab-Fab interactions are also germline-encoded (34).

The question remains whether the selection and subsequent maturation of these inter Fab contacts impacts antibody maturation or functional activity. The prevalence of an immunogenic repeat region in CSPs across different *Plasmodium* species (10), albeit with distinct repeating sequence motifs, suggests that this region may act as an immune decoy (35) by favoring selection of antibodies with Fab-Fab interactions. A study of mAbs produced by immunization with *Pf* sporozoites indicates that clonal selection of higher-affinity, germline B-cell receptors (BCRs), rather than efficient SHM, seems to drive anti-NANP responses (36). These germline BCRs may then represent the precursors of anti-NANP mAbs that display inter-Fab contacts. Consequently, it was hypothesized that high-avidity cross-linking of BCRs from homotypic contacts may signal B-cells to exit the germinal center either prematurely or with limited rounds of somatic mutation, especially in the antibody-antigen interface, and perhaps disfavor the formation of long-lived plasmablasts, which are responsible for generating high amount of circulating antibodies as immediate responses to sporozoite invasion (37). The increasing number of observations of homotypic Fab-Fab interactions in anti-NANP antibodies here and in previous studies (26, 27), and recently in an antibody targeting CSP on a different *Plasmodium* species, *P. berghei* (38), support the above hypothesis. Fab-Fab interactions boost apparent antibody affinity through avidity effects and are also shared features among protective anti-NANP mAbs. However, such contacts may not be beneficial for the formation of immune memory and antibody potency and thus durable antibody responses, and further suggests that short NANP repeat based immunogens to prevent or modulate homotypic contacts might have some advantages to consider in design of next-generation CSP-based malaria vaccines. Such immunogens date back to the 1980s when short NANP repeats [(NANP)_3_] conjugated to tetanus toxoid were tested in a human trial and induced strong anti-NANP antibody responses (39), but no booster effect due to the lack of T-cell epitopes e.g. from other non-repeat domains (40). Interestingly, recent immunization studies with a truncated CSP with only 9 NANP repeats induced lower BCR signaling in NANP-repeat-specific B-cells, stronger responses to N- and C-term epitopes, and protected more mice against mosquito bite challenge as compared to immunization with a longer CSP containing 27 NANP repeats (41). Whether this approach will lead to more durable antibody responses and robust immune memory remains to be determined.

The crystal structures reported here of additional antibodies derived from the *IGHV3-33* gene further emphasize the role of the conserved Trp^52^ in CDR H2 for interaction with Pro in the NANP repeats (Fig. 4). Alanine substitution not only of ^H^Trp^52^, but also^L^Trp^32^, and ^H^Trp^33^ in mAbs 311, 317 and 397 disrupts antibody binding considerably (29). The convergent usage of Trp is also highlighted by the interaction between ^L^Trp^96^ encoded by the light chain *J* gene in mAb366 with Asn in the NANP repeats (Fig. 5C), whereas ^L^Trp^96^ in Fab224 and 399 hydrogen bonds with the alanine backbone in the NPNA peptide (Fig. 5A, B). We also observed that ^H^Phe^50^ was present in two antibodies from the *IGHV3-49* germline for interaction with Pro in the conserved NPNA type I β-turn, similar to the Trp in *IGHV3-33* mAbs (Fig. 5A, B). Other aromatic residues, such as His and Tyr, can form van der Waals interactions with the peptide as seen in CDR H2 of *IGHV3-33* antibodies (Fig. 4C). The high prevalence of such interactions between Fab aromatic residues and the NANP peptide is summarized for anti-NANP and anti-junction mAbs in Fig. 7. A recent cryo-EM structure of mouse mAb3D11 against the repeat region of *P. berghei* CSP reveals that 3D11 uses eight aromatic residues to form an aromatic cage for antigen recognition, with a germline-encoded Tyr from the light-chain playing a key role (38). These structural insights suggest that the NANP repeats in PfCSP prime the human immune system to select antibodies from germline genes with well-positioned aromatic residues for the initial encounter. These favorable, dominant interactions with germline-encoded aromatic residues may limit SHM and represent another hurdle that the NANP repeats pose for eliciting durable and more potent human antibody responses.

**Fig 7.**
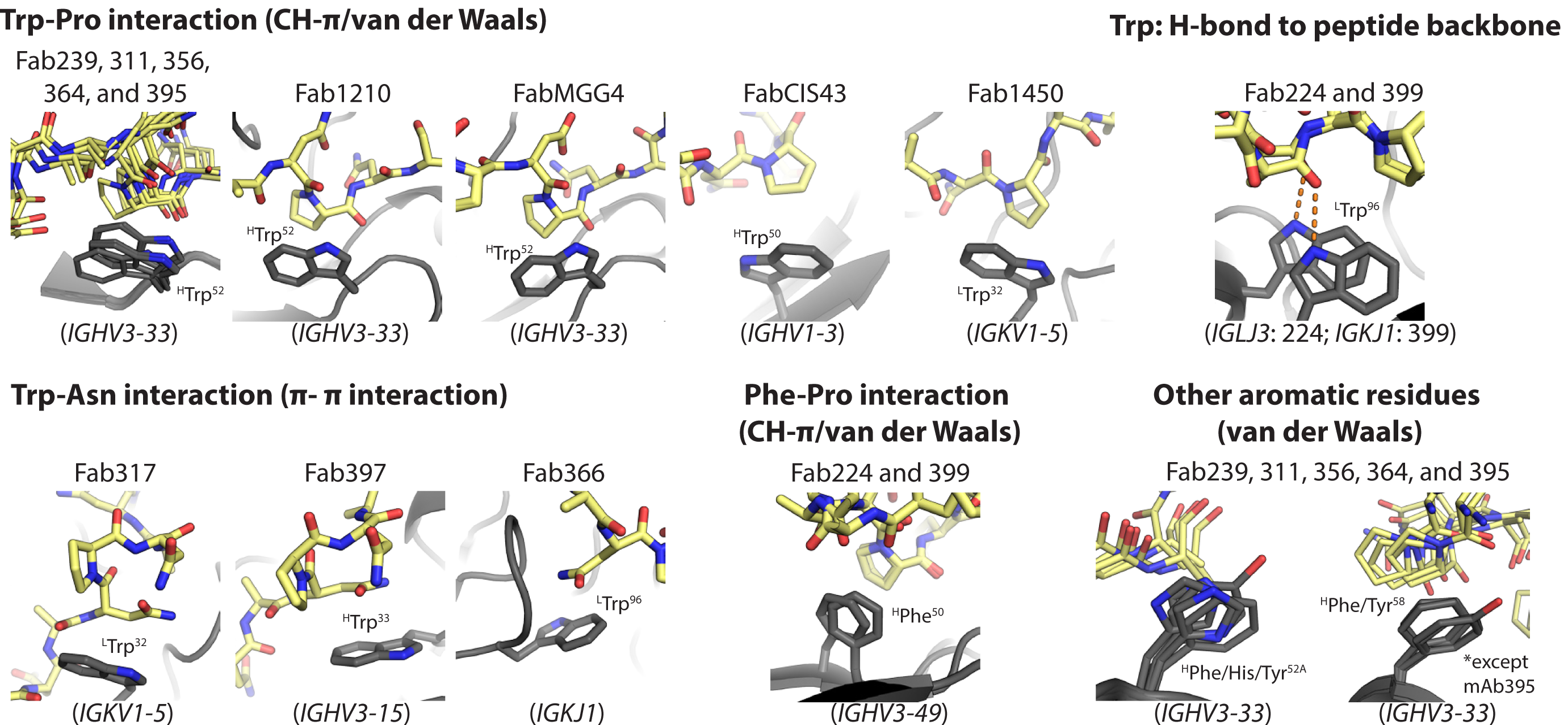
Summary of known interactions between Fab aromatic residues and the NANP or junction region. The Fab aromatic residues and CSP peptides are shown in black and yellow sticks, respectively. Hydrogen bonds are represented as orange dashes. Residue identity and number (with H and L for heavy and light chain) and the corresponding antibody germline gene is indicated. Crystal structures of Fab1210, MGG4, CIS43, 1450, and 317 and 397 were obtained from previous studies (22, 23, 26, 28, 29) (PDB ID: 6D01, 6BQB, 6B5O, 6D11, 6AXL, and 6UC5, respectively).

A recent study has shown that antibody affinity to NANP peptides correlates with inhibition of the parasite’s traversal of hepatocytes *in vitro* and also that antibodies with high affinity to NANP, rather than the other motifs (e.g. NPDP, or NVDP) in the junctional region, exhibit a high level of protection in the mouse model of parasitemia (24). Here, we also demonstrate a correlation between affinity, measured against both NANP repeats and rsCSP, and *in vivo* protection in the liver burden assay (Fig. 3, and Tables 1 and 2). The parasitemia data also follow this trend, but are not a large enough dataset to assign statistical significance. Other structural and biophysical features, which include total paratope BSA, number of hydrogen bonds between paratope and epitope, and antibody melting temperature, do not appear to correlate with *in vivo* protection (Table 2). Perhaps not surprisingly, the anti-NANP antibodies share a similar k_on_, but can differ substantially in their k_off_, which explain the lower affinities observed for mAbs 366 and 395 (Table 2). The k_off_ then dictates the linear correlation with normalized percent inhibition of parasite burden (Fig. 3). However, the caveat for the current analysis is that only two antibodies, mAbs 366 and 395, in this study have low affinity and poor *in vivo* protection, and, hence, these two data points tend to dominate the regression models. As mAbs analyzed in this study were initially screened based on their avidity on ELISA, low affinity antibodies are then likely to be underrepresented. To reduce model bias, we also performed bootstrapping to generate 1,000 models for each analysis and observed lower average R^2^ values (R^2^-boot), but that still indicate correlation with *in vivo* protection (Fig. 3). Despite these limitations, our results should serve as an important platform for development and engineering of anti-NANP mAbs, including antibody evolution using yeast display technologies.

Likewise, structural analysis reveals features on both the paratopes and epitopes that may contribute to low affinity antibody and, consequently, poor protection. One correlate that we observe is that the high affinity protective mAbs all recognize epitopes with secondary structural motifs, consisting of a type I β-turn and Asn pseudo 3_10_ turn, which represent the repeating unit of the long-range spiral form of rsCSP (Fig. 6). Conversely, low affinity, less protective antibodies possess epitopes with few to no structural motifs (Table 2, and Fig. 6). A restricted binding groove, and consequently short epitope with a single type I β-turn, likely contributes to the low affinity of Fab395 (Figs. 4B and 6). On the other hand, the low affinity and less protective Fab366 recognizes an extended conformation of NANP repeats that lack any secondary structural motifs with its shallow groove (Figs. 5C and 6). Intriguingly, the non-protective and low affinity Fab1450 (26) also shares similar features of utilizing a shallow groove to bind an extended NANP epitope.

Consequently, structural motifs such as the repeating type I β-turn and Asn pseudo 3_10_ turn as in the spiral rsCSP could be incorporated into the design of next-generation immunogens, also with shorter length designs to prevent homotypic interactions. Future studies to explore anti-NANP mAbs from different germline genes and/or immunization trials will help verify this hypothesis and/or contribute additional structural properties that influence binding affinity and *in vivo* protection. Other factors such as pharmacokinetics may impact antibody protective capacity *in vivo* and require further examination. Overall, the findings here should aid in defining the optimal characteristics of anti-NANP antibodies for therapeutic use, and also guide the design of more effective vaccines against malaria.

## Methods

### Antibody production

For protection studies, all mAbs were made as IgG1 and expressed in Chinese hamster ovary cells (ExpiCHO; Thermo Fisher Scientific, Waltham, MA). The mAbs were purified using HiTrap Protein A HP column (GE Healthcare, Chicago, IL), followed by size exclusion chromatography (Superdex 200 16/90; GE Healthcare, Chicago, IL) and washed with 0.5 M Arginine in Dulbecco’s PBS pH 7.4 (DPBS: Thermo Fisher, Waltham, MA) as described previously (42) to remove possible endotoxins. The absence of endotoxin contamination was determined using Endosafe® nexgen-PTS™ portable endotoxin testing system (Charles River, Wilmington, MA). For structural and biophysical characterizations, all Fabs were expressed in ExpiCHO cells and purified using a HiTrap Protein G HP column (GE Healthcare, Chicago, IL) followed by size exclusion chromatography as used for the IgG1 but in Tris Buffered Saline (TBS: 50 mM Tris pH 8.0, 137 mM NaCl, 3.6 mM KCl). rsCSP was expressed in *E. coli* (SHUFFLE cells; New England Biolabs, Ipswich, MA) and purified as described. All synthetic NANP peptides in this study were purchased from Innopep Inc. (San Diego, CA).

### Assessment of *in vivo* protection

Experiments were performed as described previously (32, 33). Briefly, to measure liver burden, mice (*N* = 5) were IV injected with 100 µg of mAb per mouse and, 16 h later, challenged IV with 2000 *P. berghei* transgenic sporozoites expressing the *P. falciparum* CSP and luciferase. 42 h after challenge, mice were injected IP with 100 µl of D-luciferin (30 mg/mL), having previously been anesthetized by exposure to isoflurane. Bioluminescence in the liver was measured using an IVIS Spectrum (Perkin Elmer, Waltham, MA). For the blood-stage parasitemia study, mice (*N* = 6) were passively immunized with 100 or 300 µg of mAb and, 16 h later, recipient mice and controls were anesthetized with 2% Avertin prior to challenge by a 10-minute exposure to the bites of 7 mosquitoes of which 5 on average are infected with the transgenic parasite. Parasite infection of red blood cells was assessed from day 4 after challenge by microscopic observation of blood smears. All procedures were performed according to ACUC procedures at Johns Hopkins University.

### Isothermal titration calorimetry

ITC experiments were performed on a MicroCal Auto-iTC200 (GE Healthcare, Chicago, IL). Prior to the measurement, all Fabs were extensively dialyzed against DPBS. The peptides were placed in the syringe at a concentration of ∼150 μM for Ac-NPNA NPNA-NH_2_ (NPNA_2_), ∼80 μM for Ac-NPNANPNA NPNA-NH_2_ (NPNA_3_), ∼60 μM for Ac-NPNANPNA NPNANPNA-NH_2_ (NPNA_4_), ∼40 µM for Ac-NPNANPNA NPNANPNA NPNANPNA-NH_2_ (NPNA_6_), and ∼40 μM for Ac-NPNANPNA NPNANPNA NPNANPNA NPNANPNA-NH_2_ (NPNA_8_), whereas the concentration of Fab in the cell was ∼10 μM for all experiments. The Fab and peptide concentrations were determined by UV absorbance at 280 nm and 205 nm, where the molar extinction coefficients for the peptides at 205 nm were estimated as described previously (43). The titrations were all performed with peptides in the syringe and antibodies in the cell and consisted of 16 injections of 2.45 μl peptide for experiments with NPNA_2_ and 32 injections for other experiments at a rate of 0.5 μl/s at 120 s time intervals, with injection duration of 4.9 s, injection interval of 180 s, and reference power of 5 μCal. Experiments were conducted in triplicate (*N* = 3) at 25°C. Fitting of the integrated titration peaks was performed with Origin 7.0 software using a single-site binding model. The first data point was excluded from the fit as common practice.

### Bio-layer interferometry

The binding of all Fabs to biotinylated-rsCSP was measured using bio-layer interferometry (Octet Red; Pall ForteBio, Fremont, CA). Biotinylated-rsCSP were loaded onto streptavidin biosensors (Pall ForteBio, cat No 18-5019) at 10 μg/mL in kinetics buffer (TBS + 0.002 % Tween20 and 0.01 % BSA). The loaded sensors were dipped into solutions containing dilutions of each Fab in Kinetics buffer at a concentration of 1000, 500, 250, 125, and 62.5 nM, respectively (except for Fab317, the serial dilution concentrations are 250, 125, 62.5, 31.25, 15.63 nM, respectively). The binding experiments were performed with the following steps: 1) baseline in kinetics buffer for 60 s; 2) loading of rsCSP for 60 s; 3) baseline for 60 s; 4) association of antibody for 60 s; and 5) dissociation of antibody into kinetics buffer for 120 s. A reference well with no rsCSP loaded onto the sensor was run in all experiments and subtracted from sample wells to correct for drift and buffer evaporation. Octet assays were carried out at 25 °C. Data were analyzed using the Octet Red Data Analysis software version 9.0.

### Differential scanning calorimetry

The thermal stability of all IgG1 in Dulbecco’s PBS (Thermo Fisher, Waltham, MA) from 20 to 110 °C was measured using a MicroCal VP-Capillary calorimeter (Malvern, UK) at a scanning rate of 90 °C/h. Data were analyzed using the VP-Capillary DSC automated data analysis software and fit to a non-two-state model.

### X-ray crystallography and structural analysis

Fabs 239, 356, 364, 395, 224, 250, 399, and 366 were concentrated to 10 mg/ml and mixed with either NPNA_2_, NPNA_3_, NPNA_4_, or NPNA_6_ peptide in a 1:5 molar ratio of Fab to peptide. Six substitutions and one deletion (from ^112^SSASTKG^118^ to ^112^VSRRLP^117^) were introduced into the elbow region of Fab395 and Fab366 heavy chains, and different mutations (from ^112^SSASTKG^118^ to ^112^FNQIKG^117^) were introduced to the elbow region of Fab364 heavy chain to stabilize the Fab and facilitate crystallization as previously described (44). Additionally, Fab239-NPNA_2_, Fab364-NPNA_2_ and Fab250-NPNA_3_ co-complexes were mixed with Streptococcal immunoglobulin G-binding protein G (domain III) in the Fab to protein G ratio of 1:1. Domain III of protein G has also previously been shown to enhance the crystallizability of Fabs (45). Crystal screening of Fab-peptide complexes was performed using our high-throughput, robotic CrystalMation system (Rigaku, Carlsbad, CA) at The Scripps Research Institute, using the sitting drop vapor diffusion method with a 35 μL reservoir solution and each drop consisting of 0.1 μL protein + 0.1 μL precipitant. Fab239-NPNA_2_ co-crystals were grown in 0.2 M NaCl, and 20% (w/v) PEG 3350 at 20°C as precipitant and were cryoprotected in 20% ethylene glycol. Fab356-NPNA_2_ crystals and Fab366-NPNA_3_ grew in 40% PEG-600, and 0.1M CHES pH 9.5 with final pH of 9.6 at 4°C. Fab364-NPNA_2_ crystals grew in 20% PEG-8000, and 0.05 M KH_2_PO_4_ at 4°C and were cryoprotected in 20% glycerol. Fab395-NPNA_2_ crystals grew in 30% PEG-4000, 0.2 M ammonium acetate, and 0.1 M sodium citrate pH 5.6 at 20°C. Fab224-NPNA_4_ crystals and Fab239-NPNA_4_ grew in 20% PEG-6000, and 0.1 M HEPES pH 7.0 at 20°C and were cryoprotected in 20% ethylene glycol. Fab399-NPNA_3_ crystals grew in 20% PEG 3350, and 0.2 M potassium fluoride pH 7.2 at 20°C and were cryoprotected in 20% ethylene glycol. Fab250-NPNA_4_ crystals grew in 1.6 M ammonium sulfate, and 0.1 M citric acid pH 4.0 at 20°C and were cryoprotected in 20% glycerol. Fab399-NPNA_6_ crystals grew in 50% MPD, 0.2 M ammonium dihydrogen phosphate, 0.1 M Tris pH 8.5 at 20°C. X-ray diffraction data were collected at the Advanced Proton Source (APS) beamline 23ID-B or beamline 23IDD, or at the Stanford Synchrotron Radiation Lightsource beamline 12-2, and processed and scaled using the HKL-2000 package (46). The structures were determined by molecular replacement using Phaser (47). Structure refinement was performed using phenix.refine (48) and iterations of refinement using Coot (49). Amino-acid residues of the Fabs were numbered using the Kabat system, and the structures were validated using MolProbity (50). For structural analysis, buried surface areas (BSAs) were calculated with the program MS (51), and hydrogen bonds were assessed with the program HBPLUS (52). The crystal structures of all Fab-peptide complexes have been deposited in the Protein Data Bank with accession codes: 6W00 (Fab239-NPNA_2_), 6W05 (Fab356-NPNA_2_), 6WFW (Fab364-NPNA_2_), 6WFX (Fab395-NPNA_2_), 6WFY (Fab224-NPNA_4_), 6WFZ (Fab399-NPNA_3_), 6WG0 (Fab366-NPNA_3_), 6WG1 (Fab399-NPNA_6_), and 6WG2 (Fab239-NPNA_4_).

### Statistical analysis

The parasite liver burden load data (*N = 5* mice) were compared for significance using a Mann-Whitney U test, whereas the blood-stage parasitemia data (*N = 6* mice) were analyzed using the log-rank test, where *p < 0*.*05* (*) and *p < 0*.*01* (**) indicated levels of statistically significant differences. The liver burden data were reported as the geometric mean of the total flux with 95% confidence interval (Fig. 1A). All statistical parameters for the mouse *in vivo* studies were calculated with the Hmisc (liver burden data), and the survival and survminer packages (parasitemia data), and the graphs were plotted with the ggplot2 package in R. Bootstrapping for the linear regression models was performed with the caret package and also plotted with the ggplot2 in R. Each ITC experiment was performed with three replicates (*N = 3*), and the data are reported as the arithmetic mean ± SD.

## Acknowledgements

We thank Robyn Stanfield, Xueyong Zhu, and Xiaoping Dai (The Scripps Research Institute) for advice regarding crystal structure determination, and Randall MacGill and Ashley Birkett for careful reading and comments on the manuscript. This work was supported by PATH’s Malaria Vaccine Initiative and the Bill & Melinda Gates Foundation (grant no. OPP1170236) under collaborative agreements with The Scripps Research Institute. Y.F-G. and F.Z. thank Bloomberg Philanthropies for continued support. We thank other members of the collaborative team who identified antibodies for further analysis from the Mal071 trial: Christian Ockenhouse, Ulrike Wille-Reece and Scott Gregory (PATH), Erik Jongert and Robert van den Berg (GSK), Bill Robinson (Atreca), and Sheetij Dutta, Jason Regules, Elke Bergmann-Leitner and Michele Spring (WRAIR).

