## Supplementary Material for "Structural and biophysical correlation of anti-NANP antibodies with *in vivo* protection against *P. falciparum*"

**Tossapol Pholcharee^1^, David Oyen^1^**^†^**, Yevel Flores-Garcia^2^, Gonzalo Gonzalez-Paez^1^, Zhen Han^1‡^, Katherine L. Williams^3^, Daniel Emerling^3^, Wayne Volkmuth^3^, Emily Locke^4^, C. Richter King^4^, Fidel Zavala^2^, Ian A. Wilson^1,5^***

^1^Department of Integrative Structural and Computational Biology, The Scripps Research Institute, La Jolla, CA 92037, USA.

^2^Malaria Research Institute, Johns Hopkins Bloomberg School of Public Health, Baltimore, MD 21204, USA.

^3^Atreca Inc., South San Francisco, CA 94080, USA.

^4^PATH’s Malaria Vaccine Initiative, Washington, DC 20001, USA.

^5^The Skaggs Institute for Chemical Biology, The Scripps Research Institute, La Jolla, CA 92037, USA.

^†^Current address: Pfizer Inc., San Diego, CA 92121, USA.

^‡^Current address: Wondfo USA Co., Ltd., San Diego, CA 92121, USA.

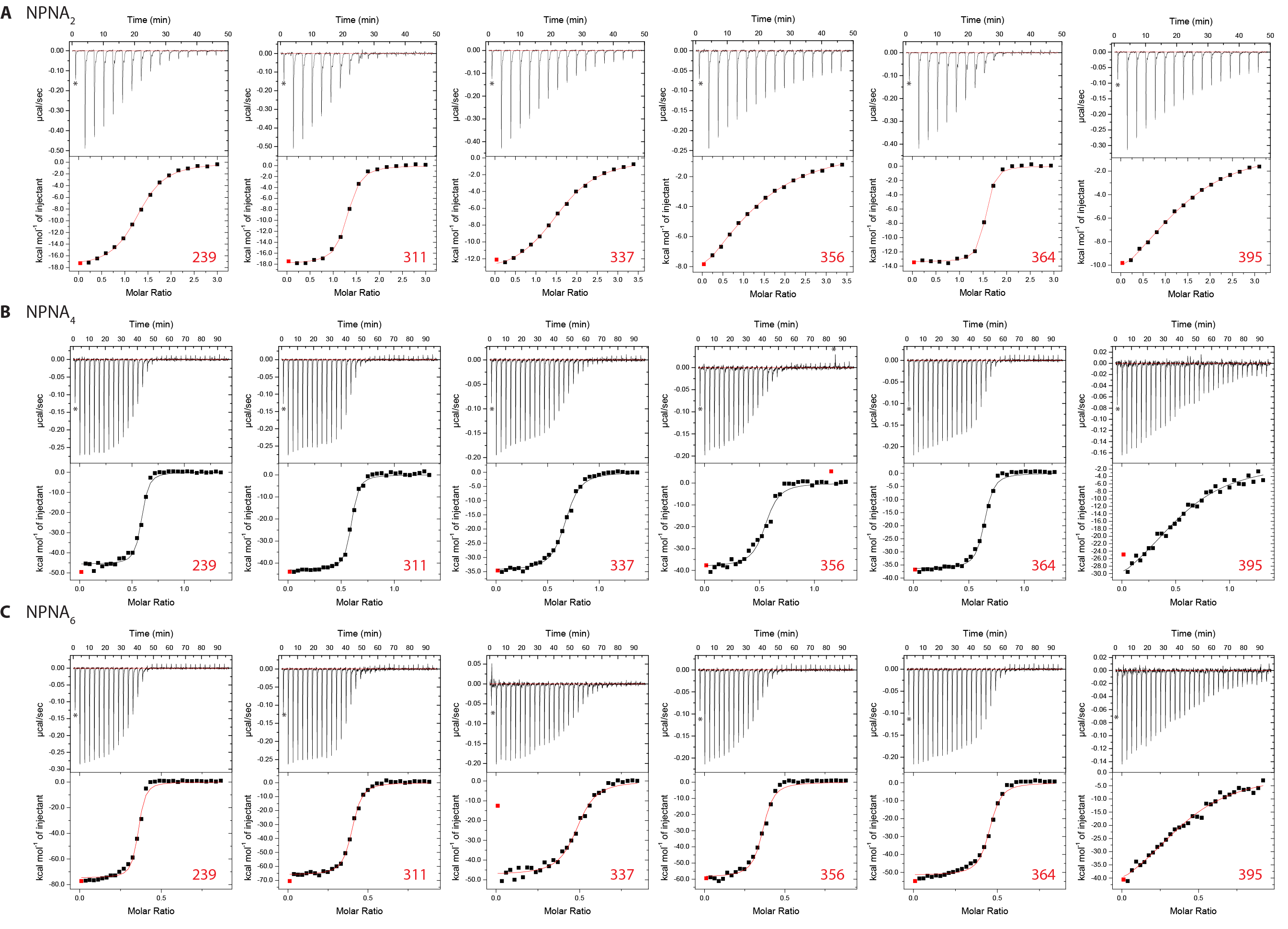

**Fig. S1. ITC binding curves for the *IGHV3-33* Fabs.** ITC binding data for each indicated Fab with: **A)** 8-mer peptide Ac-NPNANPNA-NH_2_, **B)** 16-mer peptide Ac-NPNANPNA NPNANPNA-NH_2_, and **C)** 24-mer Ac-NPNANPNA NPNANPNA NPNANPNA-NH_2_. The first data point is not included in the fit and indicated by an asterisk and red square.

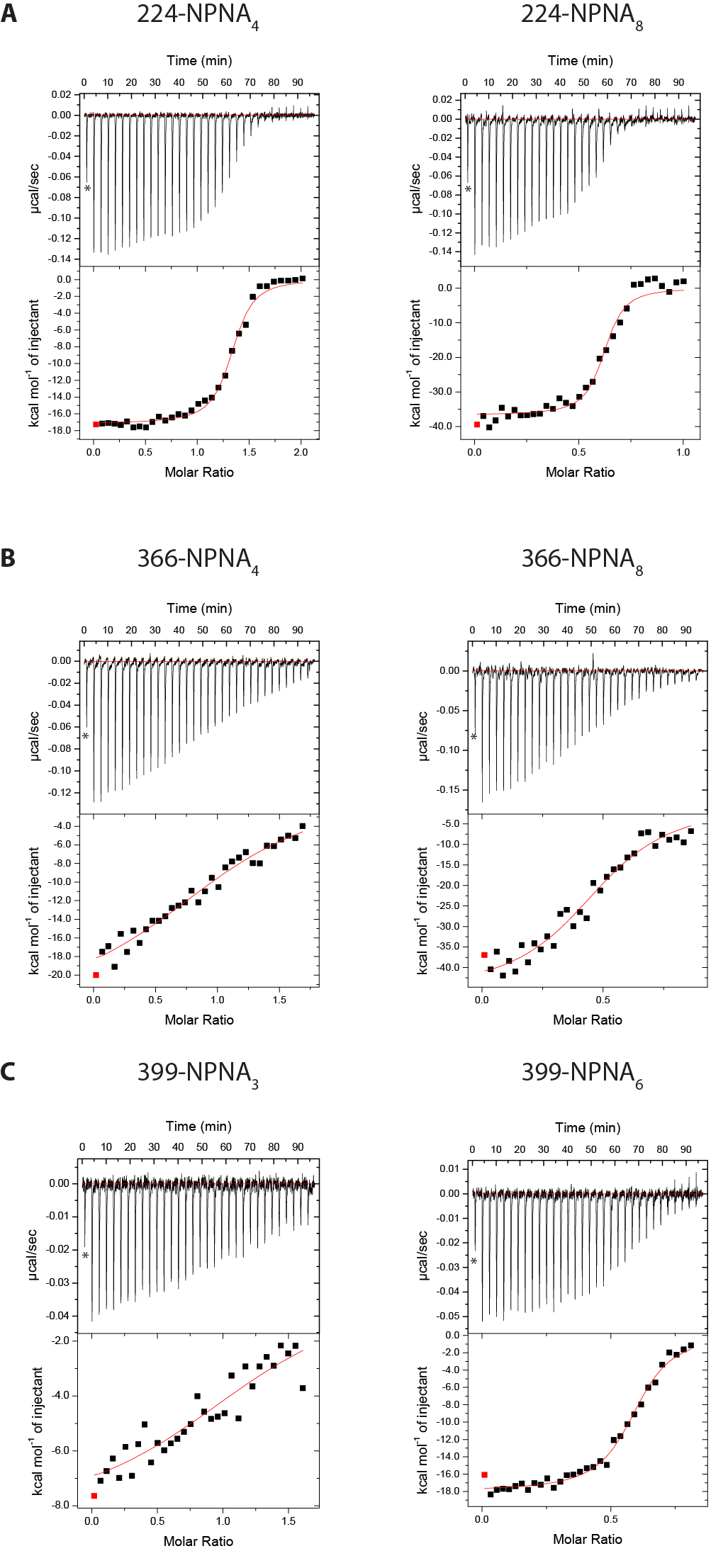

**Fig. S2. ITC binding curves for Fab224, 366, and 399 binding to NANP repeat peptides.** ITC binding data for **A)** Fab224 and **B)** Fab366 with: 1) 16-mer peptide Ac-NPNANPNA NPNANPNA-NH_2_, and 2) 32-mer Ac-NPNANPNA NPNANPNA NPNANPNA NPNANPNA-NH_2_. **C)** Fab399 was measured against: 1) 12-mer peptide Ac-NPNANPNA NPNA-NH_2_ and 2) 24-mer peptide Ac-NPNANPNA NPNANPNA NPNANPNA-NH_2_. The first data point is not included in the fit and indicated by an asterisk and a red square.

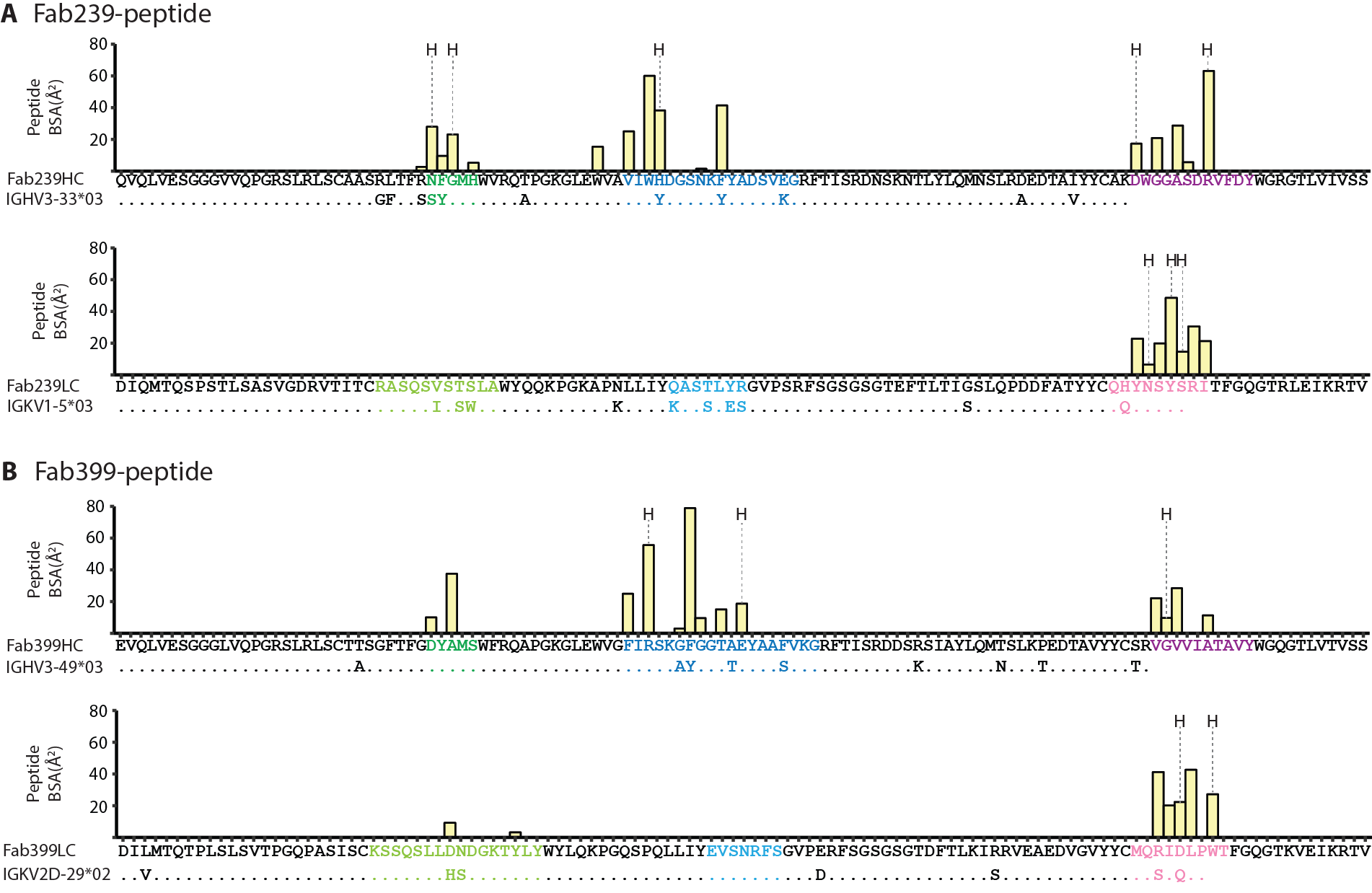

**Fig. S3. Individual residue contributions to the BSA of the Fab-peptide interface.** BSAs are shown in yellow bars for the heavy and light chains of **A)** Fab239 and **B)** Fab399. CDRs are colored as green, blue, magenta, light green, light blue, and pink for CDR H1, H2, H3, L1, L2, and L3, respectively. Additionally, the alignment between the Fab heavy/light chain sequences and germline *IGHV* and *IGKV* gene sequences also indicates somatically mutated residues. The letter “H” marks residues that are engaged in hydrogen bonds.

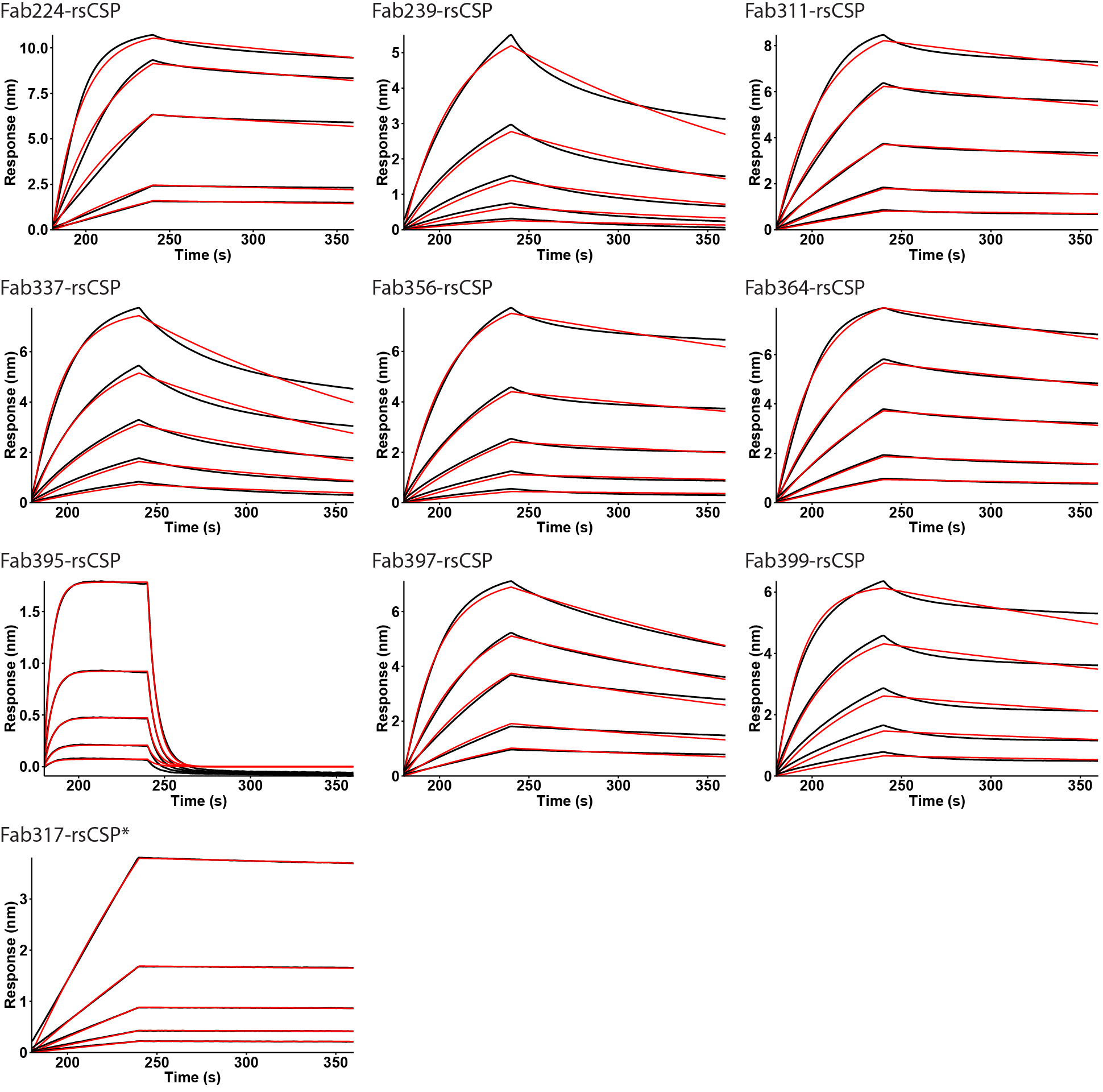

**Fig. S4. Kinetics of binding for mAbs analyzed in this study with rsCSP.** Binding was assessed using bio-layer interferometry (BLI). Binding curves are shown in black and fits are shown in red. From the top to bottom curves, Fab concentrations are 1000, 500, 250, 125, and 62.5 nM, respectively (except for Fab317 where serial dilutions are 250, 125, 62.5, 31.25, 15.63 nM, respectively).

**Table S1. The *p*-values from the pairwise comparison of mAbs in the liver burden assay using the Mann-Whitney U test (**p < 0.05,* and ***p < 0.01*).**

| **Panel 1** | **Naïve** | **311** | **239** | **337** | **356** | **364** | **395** |
| --- | --- | --- | --- | --- | --- | --- | --- |
| **Naïve** | - |  |  |  |  |  |  |
| **311** | 0.0079** | - |  |  |  |  |  |
| **239** | 0.0079** | 0.0556 | - |  |  |  |  |
| **337** | 0.0079** | 0.0079** | 0.0079** | - |  |  |  |
| **356** | 0.0079** | 0.1508 | 0.8413 | 0.0079** | - |  |  |
| **364** | 0.0079** | 0.1508 | 0.8413 | 0.0079** | 1.0000 | - |  |
| **395** | 0.0159* | 0.0079** | 0.0079** | 0.0079** | 0.0079** | 0.0079** | - |
| **Panel 2** | **Naïve** | **311** | **399** | **224** | **366** | **397** | **317** |
| **Naïve** | - |  |  |  |  |  |  |
| **311** | 0.0079** | - |  |  |  |  |  |
| **399** | 0.0079** | 0.5476 | - |  |  |  |  |
| **224** | 0.0079** | 0.4206 | 0.0556 | - |  |  |  |
| **366** | 0.0079** | 0.0952 | 0.0318* | 0.0079** | - |  |  |
| **397** | 0.0079** | 0.0952 | 0.0079** | 0.0079** | 0.6905 | - |  |
| **317** | 0.0079** | 0.6905 | 0.6905 | 0.1508 | 0.0556 | 0.0079** | - |

**Table S2. Additional isothermal titration calorimetry measurements for anti-NANP antibodies analyzed in this study.**

| **mAb** | **NPNA_2_** | | | | | **NPNA_4_** | | | | | **NPNA_6_** | | | | |
| --- | --- | --- | --- | --- | --- | --- | --- | --- | --- | --- | --- | --- | --- | --- | --- |
|  | **No.**  **of sites** | **ΔH**  **(cal/mol)** | **ΔS**  **(cal/**  **mol·K)** | **-TΔS**  **(cal/mol)** | **-ΔG**  **(cal/**  **mol)** | **No.**  **of sites** | **ΔH**  **(cal/mol)** | **ΔS**  **(cal/**  **mol·K)** | **-TΔS**  **(cal/mol)** | **-ΔG**  **(cal/**  **mol)** | **No.**  **of sites** | **ΔH**  **(cal/mol)** | **ΔS**  **(cal/**  **mol·K)** | **-TΔS**  **(cal/mol)** | **-ΔG**  **(cal/**  **mol)** |
| **Fab239** | 1.34 ± 0.01 | -17963 ± 23 | -31.9 ± 0.2 | 9511 ± 52 | -8452 ± 57 | 0.572 ± 0.011 | -46133 ± 437 | -120 ± 1.5 | 35679 ± 455 | -10454 ± 631 | 0.348 ± 0.005 | -74853 ± 179 | -214 ± 0.6 | 63903 ± 172 | -10950 ± 248 |
| **Fab311** | 1.27 ± 0.06 | -17990 ± 62 | -29.1 ± 0.3 | 8686 ± 96 | -9304 ± 114 | 0.578 ± 0.002 | -43303 ± 155 | -109 ± 1.2 | 32598 ± 144 | -10705 ± 212 | 0.386 ± 0.005 | -65990 ± 184 | -186 ± 1.0 | 55456 ± 298 | -10534 ± 350 |
| **Fab337** | 1.32 ± 0.02 | -16670 ± 87 | -29.3 ± 0.3 | 8726 ± 86 | -7944 ± 122 | 0.663 ± 0.006 | -37577 ± 5096 | -92 ± 15.3 | 27519 ± 4571 | -10058 ± 6846 | 0.480 ± 0.007 | -46407 ± 1529 | -122 ± 5.5 | 36474 ± 1642 | -9933 ± 2244 |
| **Fab356** | 1.29 ± 0.08 | -13543 ± 784 | -22.1± 2.5 | 6599 ± 757 | -6944 ± 1090 | 0.542 ± 0.006 | -38697 ± 492 | -97 ± 1.7 | 29020 ± 494 | -9677 ± 697 | 0.354 ± 0.001 | -58750 ± 95 | -163 ± 0.6 | 48499 ± 172 | -10251 ± 196 |
| **Fab364** | 1.41± 0.00 | -13877 ± 81 | -13.0 ± 0.3 | 3886 ± 75 | -9991 ± 110 | 0.634 ± 0.008 | -36087 ± 281 | -86 ± 0.9 | 25780 ± 277 | -10307 ± 395 | 0.445 ± 0.001 | -51720 ± 349 | -139 ± 1.2 | 41343 ± 344 | -10377 ± 490 |
| **Fab395** | 1.35 ± 0.02 | -14510 ± 403 | -24.8 ± 1.3 | 7404 ± 394 | -7106 ± 564 | 0.608 ± 0.032 | -36090 ± 3353 | -94 ± 11.9 | 28106 ± 3551 | -7984 ± 4884 | 0.432 ± 0.016 | -47443 ± 4643 | -132 ± 16.2 | 39256 ± 4820 | -8187 ± 6693 |
| **mAb** | **NPNA_4_** | | | | | **NPNA_8_** | | | | |  | | | |  |
|  | **No.**  **of sites** | **ΔH**  **(cal/mol)** | **ΔS**  **(cal/**  **mol·K)** | **-TΔS**  **(cal/mol)** | **-ΔG**  **(cal/**  **mol)** | **No.**  **of sites** | **ΔH**  **(cal/mol)** | **ΔS**  **(cal/**  **mol·K)** | **-TΔS**  **(cal/mol)** | **-ΔG**  **(cal/**  **mol)** |  |  |  |  |  |
| **Fab224** | 1.30 ± 0.02 | -17167 ± 67 | -24.5 ± 0.3 | 7315 ± 91 | -9852 ± 113 | 0.610 ± 0.04 | -36250 ± 552 | -87.4 ± 1.2 | 26068 ± 346 | -10182 ± 651 |  |  |  |  |  |
| **Fab366** | 1.21 ± 0.10 | -22650 ± 1274 | -50.6 ± 4.3 | 15096 ± 1291 | -7554 ± 1814 | 0.539 ± 0.028 | -41583 ± 3702 | -110.3 ± 12.7 | 32886 ± 3789 | -8697 ± 5297 |  |  |  |  |  |
| **Fab397*** | 1.14 ± 0.01 | -19915 ± 102 | -33.2 ± 0.5 | 9884 ± 147 | -10031 ± 179 | 0.560 ± 0.010 | -40090 ± 143 | -100.7 ± 0.6 | 30024 ± 179 | -10066 ± 229 |  |  |  |  |  |
| **mAb** | **NPNA_3_** | | | | | **NPNA_6_** | | | | |  | | | |  |
|  | **No.**  **of sites** | **ΔH**  **(cal/mol)** | **ΔS**  **(cal/**  **mol·K)** | **-TΔS**  **(cal/mol)** | **-ΔG**  **(cal/**  **mol)** | **No.**  **of sites** | **ΔH**  **(cal/mol)** | **ΔS**  **(cal/**  **mol·K)** | **-TΔS**  **(cal/mol)** | **-ΔG**  **(cal/**  **mol)** |  |  |  |  |  |
| **Fab399** | 1.34 ± 0.06 | -8172 ± 548 | -1.8 ± 2.1 | 543 ± 636 | -7629 ± 840 | 0.591 ± 0.002 | -17983 ± 216 | -28.1 ± 1.2 | 8378 ± 366 | **-**9605 ± 425 |  |  |  |  |  |
| **Fab317**^†^ | 1.27 ± 0.02 | −15,700 ± 225 | −20 ± 1 | 5963 ± 298 | -9737 ± 373 |  |  |  |  |  |  |  |  |  |  |

*Data obtained from (29)*.*

^†^Data obtained from (28)*.*

**Table S3. X-ray data collection and refinement statistics.**

| **Data collection** | **Fab399-NPNA_6_** | **Fab239-NPNA_4_** | **Fab366-NPNA_3_** | **Fab399-NPNA_3_** |
| --- | --- | --- | --- | --- |
| Beamline | APS23-IDB | SSRL12-2 | APS23-IDD | APS23-IDD |
| Wavelength (Å) | 1.03316 | 0.97946 | 1.03320 | 1.03324 |
| Space group | P2_1_ | P2_1_ | P2_1_2_1_2_1_ | P2_1_ |
| Unit cell parameters (Å, °) | a=62.97, b=85.90, c=89.46 | a=82.16, b=55.70, c=115.73 | a=59.66, b=68.71, c=108.79 | a=63.40, b=87.60, c=89.65 |
|  | α=90, β=100.3, γ=90 | α=90, β=98.9, γ=90 | α=β=γ=90 | α=90, β=101.0, γ=90 |
| Resolution (Å) | 50.00-2.10 (2.14-2.10)^a^ | 50.00-2.54 (2.58-2.54)^a^ | 50.00-1.60 (1.63-1.60)^a^ | 50.00-1.85 (1.88-1.85)^a^ |
| Unique Reflections | 49,542 (1,393)^a^ | 31,852 (1,613)^a^ | 56,859 (1,420)^a^ | 80,717 (3,088)^a^ |
| Redundancy | 2.4 (1.8)^a^ | 3.7 (3.6)^a^ | 11.7 (2.9)^a^ | 3.0 (1.9)^a^ |
| Completeness (%) | 89.1 (51.0)^a^ | 91.9 (93.6)^a^ | 95.1 (48.4)^a^ | 97.2 (74.4)^a^ |
| <I/σ_I_> | 10.3 (1.2)^a^ | 7.9 (1.6)^a^ | 28.9 (1.0)^a^ | 18.6 (1.1)^a^ |
| R_sym_^b^ (%) | 8.2 (40.7)^a^ | 14.5 (89.8)^a^ | 7.6 (55.9)^a^ | 7.9 (48.4)^a^ |
| R_pim_^b^ (%) | 6.0 (33.3)^a^ | 8.2 (52.2)^a^ | 2.2 (32.6)^a^ | 5.2 (38.0)^a^ |
| CC_1/2_^c^ (%) | 90.1 (71.5)^a^ | 88.0 (52.1)^a^ | 95.4 (71.9)^a^ | 91.7 (69.2)^a^ |
| **Refinement statistics** |  |  |  |  |
| Resolution (Å) | 44.01-2.10 | 32.81-2.54 | 45.05-1.60 | 46.75-1.85 |
| Reflections (work) | 49,513 | 31,820 | 56,784 | 80,521 |
| Reflections (test) | 2,434 | 1,591 | 2,793 | 3,869 |
| R_cryst_^d^ / R_free_^e^ (%) | 19.6/24.6 | 19.5/24.6 | 18.9/21.1 | 18.6/22.5 |
| **No. of atoms** |  |  |  |  |
| Fab | 6,502 | 6,475 | 3,325 | 6,645 |
| Peptide | 168 | 115 | 79 | 158 |
| Water | 347 | 106 | 265 | 367 |
| **Average B-value (Å^2^)** |  |  |  |  |
| Fab | 37 | 42 | 24 | 39 |
| Peptide | 44 | 36 | 31 | 41 |
| Water | 38 | 36 | 33 | 42 |
| Wilson B-value | 32 | 36 | 22 | 29 |
| **RMSD from ideal geometry** |  |  |  |  |
| Bond length (Å) | 0.005 | 0.002 | 0.007 | 0.007 |
| Bond angle (°) | 0.73 | 0.58 | 0.90 | 0.85 |
| **Ramachandran statistics^f^** |  |  |  |  |
| Favored (%) | 98.17 | 97.34 | 97.73 | 98.17 |
| Outliers (%) | 0.11 | 0.00 | 0.00 | 0.11 |

^a^ Numbers in parentheses refer to the highest resolution shell.

^b^ *R*_sym_ = Σ*_hkl_* Σ*_i_* | I*_hkl,i_* - <I*_hkl_*> | / Σ*_hkl_* Σ*_i_* I*_hkl,i_* and R*_pim_* = Σ*_hkl_* (1/(n-1))^1/2^ Σ*_i_* | I*_hkl,i_* - <I*_hkl_*> | / Σ*_hkl_* Σ*_i_* I*_hkl,i_*, where I*_hkl,i_* is the scaled intensity of the i^th^ measurement of reflection h, k, l, <I*_hkl_*> is the average intensity for that reflection, and *n* is the redundancy.

^c^ CC_1/2_ = Pearson correlation coefficient between two random half datasets.

*^d^ R*_cryst_ = Σ*_hkl_* | *F*_o_ - *F*_c_ | / Σ*_hkl_* | *F*_o_ | x 100, where *F*_o_ and *F*_c_ are the observed and calculated structure factors, respectively.

^e^ *R*_free_ was calculated as for *R*_cryst_, but on a test set comprising 5% of the data excluded from refinement.

^f^ From MolProbity (50).

**Table S4. X-ray data collection and refinement statistics (continued).**

| **Data collection** | **Fab239-NPNA_2_** | **Fab356-NPNA_2_** | **Fab364-NPNA_2_** | **Fab395-NPNA_2_** | **Fab224-NPNA_4_** |
| --- | --- | --- | --- | --- | --- |
| Beamline | APS23-IDD | APS23-IDD | APS23-IDD | APS23-IDD | APS23-IDD |
| Wavelength (Å) | 1.03321 | 1.03321 | 1.03320 | 1.03324 | 1.03324 |
| Space group | P4_3_2_1_2 | P2_1_2_1_2_1_ | C222_1_ | P2_1_2_1_2_1_ | P2_1_ |
| Unit cell parameters (Å, °) | a= b=121.52, c=82.61 | a=64.14, b=81.76, c=84.05 | a=80.44, b=117.05, c=116.91 | a=60.42, b=78.66, c=90.67 | a=42.92, b=67.34, c=84.77 |
|  | α= β=γ=90 | α=β= γ=90 | α=β=γ=90 | α=β=γ=90 | α=β=γ=90 |
| Resolution (Å) | 50.00-1.85 (1.88-1.85)^a^ | 50.00-2.52 (2.56-2.52)^a^ | 50.00-2.10 (2.14-2.10)^a^ | 50.00-2.60 (2.64-2.60)^a^ | 50.00-1.23 (1.25-1.23)^a^ |
| Unique Reflections | 52,998 (2,623)^a^ | 15,560 (773)^a^ | 28,464 (556)^a^ | 13,944 (675)^a^ | 114,397 (855)^a^ |
| Redundancy | 13.3 (9.8)^a^ | 7.0 (6.2)^a^ | 5.3 (1.8)^a^ | 6.7 (6.9)^a^ | 2.9 (1.0)^a^ |
| Completeness (%) | 100.0 (100.0)^a^ | 100.0 (100.0)^a^ | 86.3 (34.8)^a^ | 99.9 (100.0)^a^ | 81.7 (12.4)^a^ |
| <I/σ_I_> | 18.6 (1.9)^a^ | 13.6 (1.6)^a^ | 13.3 (2.0)^a^ | 8.0 (2.0)^a^ | 19.1 (1.4)^a^ |
| R_sym_^b^ (%) | 18.8 (93.3)^a^ | 14.4 (105.3)^a^ | 10.8 (40.0)^a^ | 21.6 (124.4)^a^ | 5.4 (35.7)^a^ |
| R_pim_^b^ (%) | 5.3 (30.3)^a^ | 5.8 (45.5)^a^ | 4.8 (31.3)^a^ | 9.0 (52.3)^a^ | 3.4 (35.3)^a^ |
| CC_1/2_^c^ (%) | 95.1 (72.5)^a^ | 90.0 (54.7)^a^ | 93.4 (73.8)^a^ | 88.3 (49.6)^a^ | 89.3 (67.4)^a^ |
| **Refinement statistics** |  |  |  |  |  |
| Resolution (Å) | 48.97-1.85 | 43.27-2.52 | 43.85-2.10 | 47.92-2.60 | 42.68-1.23 |
| Reflections (work) | 52,931 | 15,512 | 28,419 | 13,902 | 114,353 |
| Reflections (test) | 2577 | 748 | 1,431 | 699 | 5,705 |
| R_cryst_^d^ / R_free_^e^ (%) | 18.7/20.6 | 20.9/25.6 | 21.3/25.6 | 19.8/24.4 | 16.9/18.3 |
| **No. of atoms** |  |  |  |  |  |
| Fab | 6,499 | 3,281 | 3,135 | 3,202 | 3,355 |
| Peptide | 107 | 56 | 56 | 43 | 84 |
| Protein G | 882 | N/A | 453 | N/A | N/A |
| Water | 349 | 22 | 123 | 60 | 491 |
| **Average B-value (Å^2^)** |  |  |  |  |  |
| Fab | 24 | 56 | 53 | 41 | 18 |
| Peptide | 17 | 75 | 78 | 45 | 17 |
| Protein G | 26 | N/A | 45 | N/A | N/A |
| Water | 28 | 46 | 42 | 30 | 30 |
| Wilson B-value | 20 | 48 | 37 | 37 | 11 |
| **RMSD from ideal geometry** |  |  |  |  |  |
| Bond length (Å) | 0.009 | 0.004 | 0.004 | 0.002 | 0.006 |
| Bond angle (°) | 0.87 | 0.66 | 0.67 | 0.60 | 0.85 |
| **Ramachandran statistics^f^** |  |  |  |  |  |
| Favored (%) | 97.79 | 96.58 | 95.18 | 95.83 | 97.77 |
| Outliers (%) | 0.00 | 0.00 | 0.21 | 0.23 | 0.00 |

^a^ Numbers in parentheses refer to the highest resolution shell.

^b^ *R*_sym_ = Σ*_hkl_* Σ*_i_* | I*_hkl,i_* - <I*_hkl_*> | / Σ*_hkl_* Σ*_i_* I*_hkl,i_* and R*_pim_* = Σ*_hkl_* (1/(n-1))^1/2^ Σ*_i_* | I*_hkl,i_* - <I*_hkl_*> | / Σ*_hkl_* Σ*_i_* I*_hkl,i_*, where I*_hkl,i_* is the scaled intensity of the i^th^ measurement of reflection h, k, l, <I*_hkl_*> is the average intensity for that reflection, and *n* is the redundancy.

^c^ CC_1/2_ = Pearson correlation coefficient between two random half datasets.

*^d^ R*_cryst_ = Σ*_hkl_* | *F*_o_ - *F*_c_ | / Σ*_hkl_* | *F*_o_ | x 100, where *F*_o_ and *F*_c_ are the observed and calculated structure factors, respectively.

^e^ *R*_free_ was calculated as for *R*_cryst_, but on a test set comprising 5% of the data excluded from refinement.

^f^ From MolProbity (50)*.*

**Table S5. Germline genes and buried surface area (BSA) of mAbs analyzed in this study.**

| **mAb** | **HC *V* gene**  **(IGVH)** | **HC *D* gene** | **HC *J* gene** | **LC *V* gene**  **(IGKV, IGLV)** | **LC *J* gene** | **HC BSA (Å^2^)** | **LC BSA (Å^2^)** | **Total BSA**  **(Å^2^)** |
| --- | --- | --- | --- | --- | --- | --- | --- | --- |
| 239 | *3-33*03 or*04* | *D2-21*02* | *J5*01* | *KV1-5*01 or*  *KV1-5*03* | *KJ2*02 or KJ5*01* | 386 | 164 | 550 |
| 311 | *3-33*01 or *06* | *D3-22*01* | *J4*02 or J4*03* | *LV1-40*01 or*  *LV1-40*02* | *LJ3*02* | 366 | 107 | 473 |
| 337 | *3-33*03,*  *3-30*04,*14 or *16*  *3-30-3*03* | *D2-21*01* | *J3*02* | *KV3-15*01* | *KJ1*01* | N/A | N/A | N/A |
| 356 | *3-33*03* | *D3-22*01* | *J4*02 or J4*03* | *KV3-15*01* | *KJ3*01* | 457 | 115 | 572 |
| 364 | *3-33*03,*  *3-30*02, or*  *3-30-5*02* | *D4-17*01* | *J4*02* | *KV1-5*03* | *KJ1*01* | 344 | 99 | 443 |
| 395 | *3-33*03* | *D3-9*01* | *J4*02 or J4*03* | *KV1-5*01* | *KJ1*01* | 379 | 41 | 420 |
| 224 | *3-49*04* | *D4-17*01* | *J6*01* | *LV1-40*01* | *LJ3*02* | 429 | 188 | 617 |
| 399 | *3-49*03* | *-* | *J2*01* | *KV2-29*02 or KV2D-29*02* | *KJ1*01* | 314 | 178 | 492 |
| 366 | *1-2*02* | *D2-21*01* | *J3*02* | *KV1-27*01* | *KJ1*01* | 348 | 228 | 576 |
| 317  (*VH3-30*) | *3-30*14 or *16* | *D3-16*01* | *IJ4*02* | *KV1-5*01* | *KJ1*01* | 310 | 209 | 519 |
| 397  (*VH3-15*) | *3-15*01* | *D3-3*01* | *J4*02 or J4*03* | *KV2-28*01 or KV2D-28*01* | *KJ1*01, KJ2*01 or KJ5*01* | 264 | 329 | 593 |

**Table S6. Conserved residues of *IGHV3-33* mAbs that interact with the NPNA peptide**

| **mAb** | **227** | **239** | **311** | **337**† | **356** | **364** | **395** | **Germline**  **residue** | **Interaction** |
| --- | --- | --- | --- | --- | --- | --- | --- | --- | --- |
| H 31 (32)‡ | A | N | N | **T** | N* | G | C | S | Conserved MHB |
| H 32 (33)‡ | F | F | Y | **Y** | F | Y | Y | Y | Conserved vdW |
| H 33 (34)‡ | G | G | G | **G** | G | G | G | G | Conserved MHB |
| H 50 | V | V | **I** | **L** | V | **I** | V | V | **Evolved vdW** |
| H 52 | W | W | W | **W** | W | W | W | W | Conserved CH-pi |
| H 52A | Y | H | Y | **H** | H | F | H  (also SHB) | Y | Conserved MHB |
| H 58 | Y | F | F | **F** | F | Y | H | Y | Conserved vdW |
| H 95 | V | D  (SHB) | A  (MHB) | **D** | D  (SHB) | V | A  (MHB) | - | Varied as indicated |

*The main chain of ^H^Asn^31^ in mAb356 does not form a hydrogen bond with the peptide.

†Residues in 337 are included for a complete comparison but are in grey as no structure of this mAb is available.

‡Residue number for mAb395.

MHB = Hydrogen bond between the main chain and NANP peptide.

SHB = Hydrogen bond between the side chain and NANP peptide.

vdW = van der Waals interaction.
